# Nucleotide dependency analysis of DNA language models reveals genomic functional elements

**DOI:** 10.1101/2024.07.27.605418

**Authors:** Pedro Tomaz da Silva, Alexander Karollus, Johannes Hingerl, Gihanna Galindez, Nils Wagner, Xavier Hernandez-Alias, Danny Incarnato, Julien Gagneur

## Abstract

Deciphering how nucleotides in genomes encode regulatory instructions and molecular machines is a long-standing goal in biology. DNA language models (LMs) implicitly capture functional elements and their organization from genomic sequences alone by modeling probabilities of each nucleotide given its sequence context. However, using DNA LMs for discovering functional genomic elements has been challenging due to the lack of interpretable methods. Here, we introduce nucleotide dependencies which quantify how nucleotide substitutions at one genomic position affect the probabilities of nucleotides at other positions. We generated genome-wide maps of pairwise nucleotide dependencies within kilobase ranges for animal, fungal, and bacterial species. We show that nucleotide dependencies indicate deleteriousness of human genetic variants more effectively than sequence alignment and DNA LM reconstruction. Regulatory elements appear as dense blocks in dependency maps, enabling the systematic identification of transcription factor binding sites as accurately as models trained on experimental binding data. Nucleotide dependencies also highlight bases in contact within RNA structures, including pseudoknots and tertiary structure contacts, with remarkable accuracy. This led to the discovery of four novel, experimentally validated RNA structures in *Escherichia coli*. Finally, using dependency maps, we reveal critical limitations of several DNA LM architectures and training sequence selection strategies by benchmarking and visual diagnosis. Altogether, nucleotide dependency analysis opens a new avenue for discovering and studying functional elements and their interactions in genomes.

## Introduction

The basic blueprint of every living organism is encoded in its genome. High-throughput sequencing techniques have made it possible to read the genome, however, interpretation - decoding the information present in the sequence and determining its biological meaning - remains a key challenge of modern genetics. Sequence comparison across and within species is a well-established technique of genome interpretation ^1^, wherein purifying selection acting on sequence and the evolutionary relationships between genomes are leveraged to identify functional sequence elements. Sequence comparison not only leverages nucleotide-level conservation, but also statistical dependencies between nucleotides. Specifically, if the presence of a particular nucleotide at one position is associated with the presence of a particular nucleotide at another position, this suggests that these nucleotides co-evolve and thus cooperate to encode a functional element. Methods employing this principle of covariation have enabled important discoveries, particularly in protein and RNA structure biology ^2,3^. However, measuring these dependencies so far essentially relied on sequence alignment at single base-pair resolution, which can only be computed for highly conserved regions of the genome.

DNA language models (DNA LM) have recently been proposed as a technique to leverage sequence comparison without requiring alignments ^4,5^. A variety of DNA LMs have been developed, but one feature shared by all is that they are trained to predict nucleotides given their sequence context. This objective allows these models to capture evolutionary favored sequence elements and their arrangements directly from large collections of genomic sequences, without requiring additional experiments ^4^.

We and others have recently analyzed the nucleotide predictions made by DNA LMs and shown that they carry biologically meaningful information ^4–6^. For example, we found that in-vivo functional transcription factor binding motifs are generally better reconstructed by DNA LMs than non-functional copies of the same motif. Moreover, Benegas et al. found that DNA LMs predict genetic variants with phenotypic impact to be less likely than neutral variants. Several other studies have also demonstrated the value of DNA LMs to serve as so-called foundation models ^4,7–15^. In this line of research, the pre-trained DNA LMs are used as starting points for training supervised models aimed at predicting molecular phenotypes such as gene expression. In a number of cases, DNA LM-based predictors have been shown to outperform alternative approaches ^16^. These analyses indicate that DNA LMs intrinsically represent genomic functional elements. However, the foundation model paradigm employs DNA LMs as intermediate black boxes and does not reveal these elements.

In this work, we leverage DNA LMs to provide a measure of dependencies between pairs of nucleotides. We systematically study the resulting nucleotide dependency maps to determine which genomic elements they encode and exploit them to characterize functional elements and their interactions. Furthermore, we show how nucleotide dependencies can be used to compare existing DNA LM and identify their shortcomings.

## Results

### Nucleotide Dependency Maps

DNA language models are trained to reconstruct nucleotides, thereby providing nucleotide probabilities given their surrounding sequence context (Fig 1A). In principle, success at reconstructing nucleotides requires detecting characteristic genomic features more likely to be found in the sequence context. For example, the probability of a particular nucleotide in the human genome to be a guanine strongly depends on whether it is intronic (∼22%,^21^) or located at the third base of a start codon (∼100%). To study the relationship between nucleotides and their context using DNA LMs, we use a technique analogous to *in silico* mutagenesis (explained in ^22^). Specifically, we mutate a nucleotide in the sequence context (query nucleotide) into all three possible alternatives and record the change in predicted probabilities at a target nucleotide in terms of odds ratios (Fig 1B, Methods). This procedure, which can be repeated for all possible query-target combinations, quantifies the extent to which the language model prediction of the target nucleotide depends on the query nucleotide, all else equal.

**Fig. 1.**
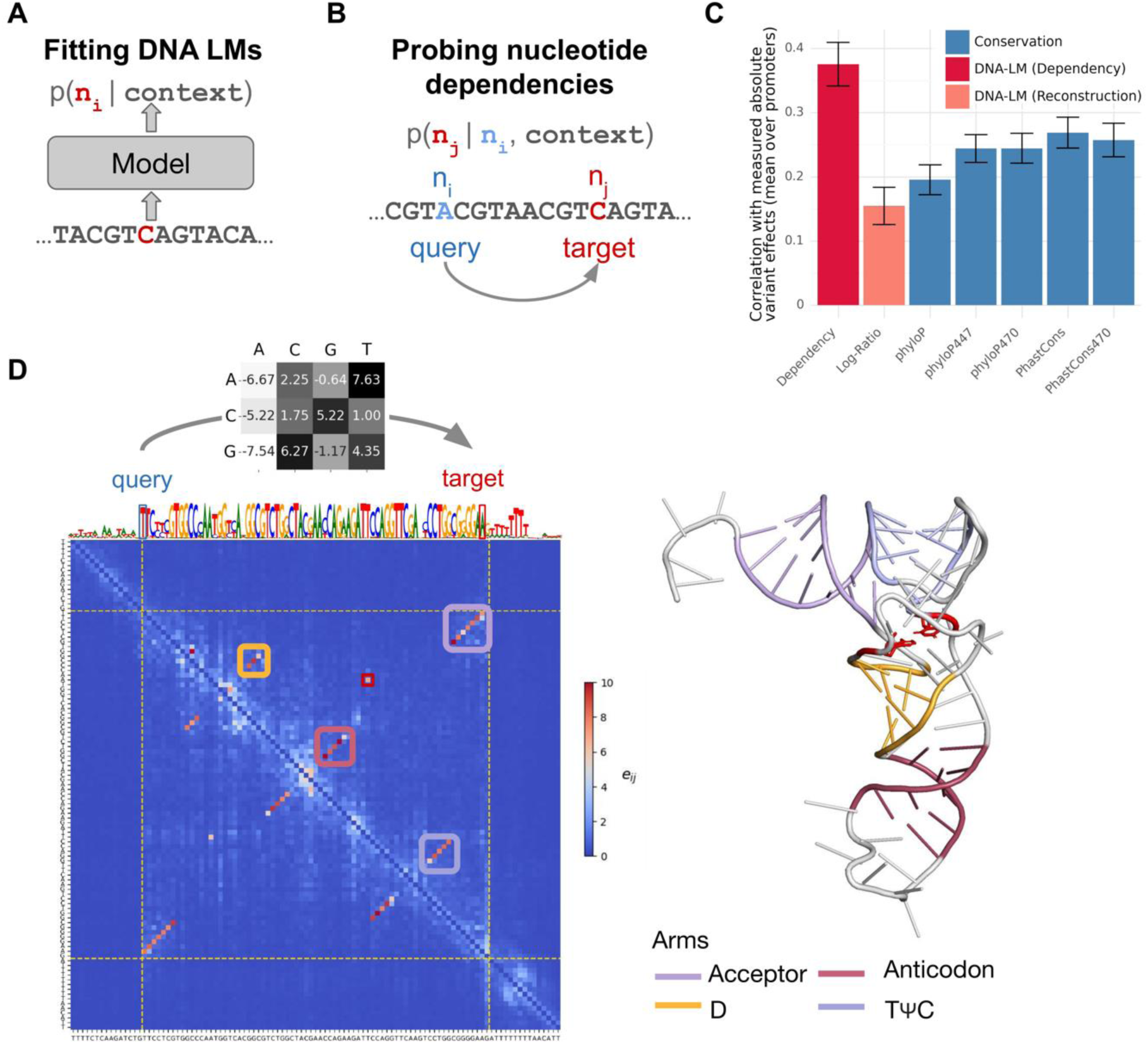
Probing nucleotide dependencies from DNA language models. **A**) DNA LMs are trained on genomes to predict nucleotides given their sequence context, assigning a probability to each one of A,C,G or T. **B**) We probe pairwise nucleotide dependencies from DNA LMs by quantifying how substituting a nucleotide at a query position affects predicted probabilities at a target position. **C)** Correlation between the absolute variant effect on gene expression, as measured using a saturation mutagenesis assay of nine human promoters (n = 8,635 variants), and the variant influence score, the reconstruction score (log-likelihood ratio of substituting the reference to the alternative nucleotide according to the DNA LM), and alignment-based conservation scores from PhyloP and PhastCons based on the 100-way, 447-way and 470-way alignment. DNA LM log ratio quantifies the reconstruction at a specific position (as depicted in **A**), while the variant influence score quantifies how each variant affects the predicted probabilities across all target nucleotides in a sequence (this study, as depicted in **B**). Error bars represent ±2 standard deviations, constructed using 100 bootstrap samples per promoter. **D**) Left. Annotated nucleotide dependency map for the *S. cerevisiae* arginine tRNA, tR(ACG)O. The grey heatmap (top) shows log-odds-ratios for all four nucleotides of a target (columns) when substituting the query nucleotide to each of the three alternatives (rows). This data is shown for the query being nucleotide 1 of the tRNA (T) and target being nucleotide 72 (A). The DNA LM log-odds ratios are consistent with the fact that these two bases encode a Watson-Crick contact in the RNA fold. The maximum absolute log-odds ratio, which defines the dependency score between those two positions, is realized when substituting an A on the query and having a T at the target. The dependency map (blue-to-red heatmap) shows dependency scores for all queries (rows) and targets (columns) in this locus. The colored rectangles in the dependency map highlight anti-parallel dependencies belonging to each of the tRNA arms while the red square delineates a dependency between 2 bases in different loops of the tRNA contributing to its tertiary structure (red bases in the tertiary structure). The track above the dependency map displays the nucleotide reconstruction predicted by the DNA LM. Right. Annotated tertiary structure of tR(ACG)O^17^.

We applied this general procedure to 14 DNA language models (STable1, Methods). Unless stated otherwise, we present results from our SpeciesLM DNA language models which were trained on regions 5’ of start codons in Fungi (SpeciesLM Fungi) and Metazoa (SpeciesLM Metazoa) (^4^, Methods). On select biological applications, we turn to other DNA language models.

As a first assessment of the biological relevance of these dependencies, we sought to verify that single nucleotide variants of known functional importance have a greater impact on DNA- LM predictions. As an aggregate score of query variant impact, we computed the average across all targets of the maximum absolute odds ratio over all possible alternative nucleotides at a target (Methods). We named this metric the variant influence score. In the ClinVar database ^23^, the influence score was significantly higher for non-coding pathogenic variants, which mostly comprised variants within or close to transcripts, than benign variants (Fig S1A, Fig S1F). This is despite using a DNA LM trained only on 2-kb regions 5’ of start codons, which only overlap a small fraction of all transcribed bases. Previous work on DNA LMs proposed to leverage reconstruction probabilities to prioritize functional variants, assuming that the less likely a variant is compared to the reference allele, the more deleterious it is ^5,6^ (Methods). Remarkably, this reconstruction-based metric showed a significantly lower performance than the influence score. However, the influence score did not outperform alignment-based scores, perhaps because criteria used for ClinVar to categorize variants as pathogenic include bioinformatics predictions which often integrate alignment-based conservation.

To ensure a less biased comparison, we next focused on a dataset from a saturation mutagenesis experiment on nine selected human promoters (Fig 1C, ^24^). Here, the variant influence score correlated with variant effect on absolute gene expression fold-change, outperforming reconstruction, as well as alignment-based conservation scores ^25–28^. Furthermore, the variant influence score also outperformed reconstruction and alignment-based conservation at distinguishing fine-mapped promoter eQTLs SNPs from matched controls, in both human and yeast (Fig S1B-E ^18,20,29,30^).

Having shown that aggregate dependency strengths reflect overall functional importance, we next sought to study them down to the level of individual query-target pairs. For every query-target pair, we considered the maximum effect a query nucleotide change has on the predicted odds of a target, yielding a 2D nucleotide dependency map (Methods). An example of such a map for the yeast arginine tRNA is given in Fig 1D. The structure of tRNAs is highly constrained, as they must fit into the translation sites of the ribosome. The entire secondary structure of the tRNA, which is defined by base pairing within the four arms of the tRNA clearly stands out with high dependencies. The dependency map also highlighted one of the tertiary structure contacts. Inspecting the underlying nucleotide predictions reveals that upon introducing single nucleotide substitutions in these pairs, the DNA-LM adapts its predictions according to Watson-Crick and, with a lesser preference, to wobble base-pairing (Fig 1D for one example). Remarkably, the RNA base-pairing rules were not a priori supplied to the model and thus were learned as a consequence of the reconstruction objective. Moreover, this structural information was captured without specifically focusing the training on tRNAs and in an alignment-free fashion.

Nucleotide pairs have two dependencies depending on which nucleotide is the query. Scoring nucleotide pairs by the maximum of those two values yielded near-perfect secondary structure contact predictions across 172 *S. cerevisiae* tRNAs. We explored alternative metrics including gradient-based dependencies and using masking instead of nucleotide substitution on query which all showed lower predictive signal; a trend that was confirmed when further assessing the dependencies on cognate donor and acceptor splice sites (Supplementary fig S1G,H).

In the following sections, we explore and categorize patterns found in nucleotide dependency maps, associate them to biological mechanisms, and exploit them to detect and characterize functional elements in the genome.

### Blocks along the diagonal highlight regulatory sequence motif instances

Within distances of a few tens of nucleotides, we noticed that sets of contiguous nucleotides often showed strong reciprocal dependencies, reflected as dense blocks along the diagonal. Many dense blocks occurred at transcription factor (TF) motif instances in promoters (Fig 2A, B). This was in striking contrast to other well reconstructed locations including repeats such as poly(dA:dT) stretches. Intuitively, all bases of a TF motif strongly depend on each other because mutations at any position could abolish or greatly reduce TF binding, thus effectively disrupting the function of the site as a whole. Therefore, we reasoned that TF motifs could be more accurately detected using DNA LMs by searching for dependency blocks, rather than using reconstruction as previously proposed ^4^.

**Fig. 2.**
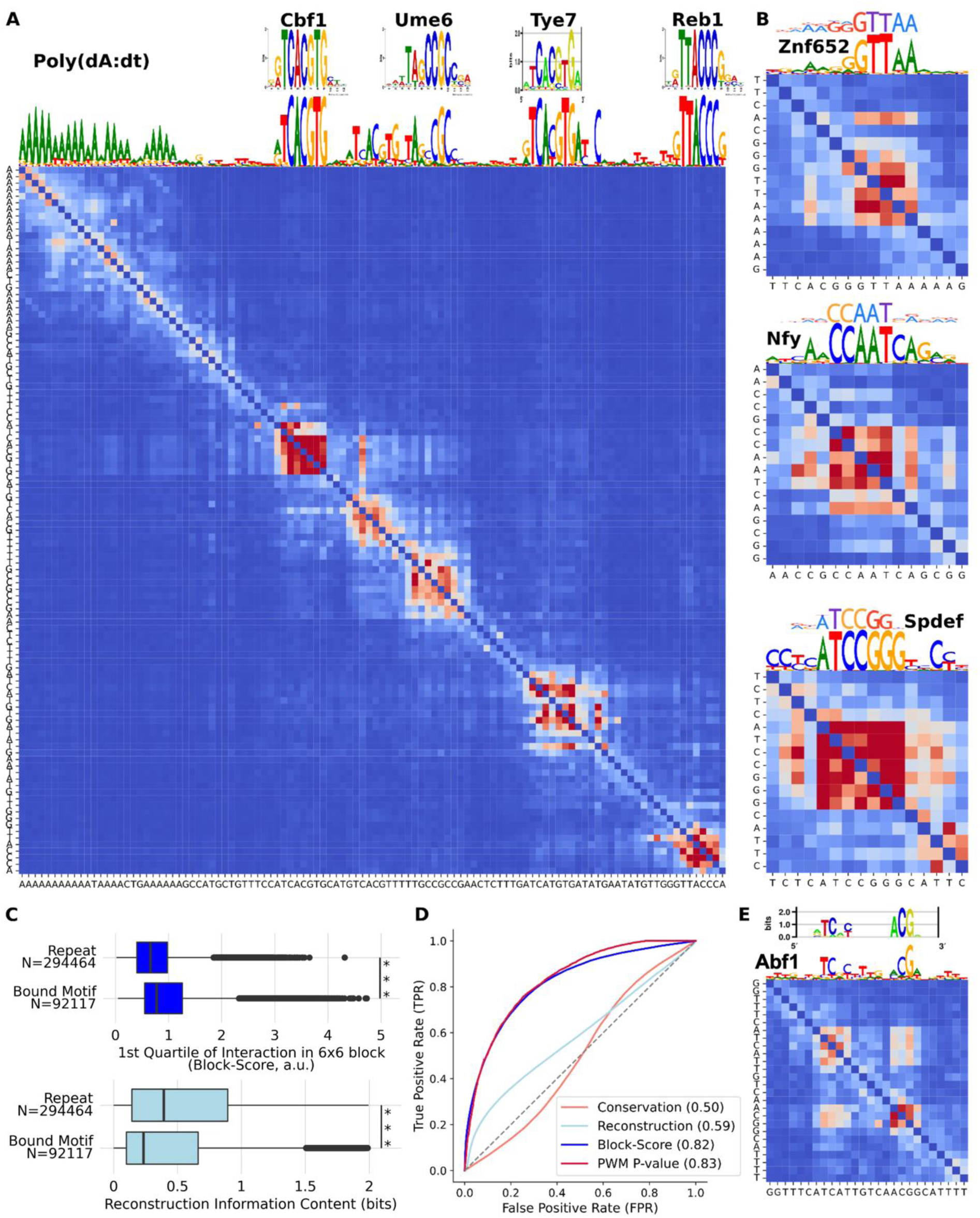
Blocks along the diagonal of dependency maps highlight regulatory sequence motif instances. **A**) SpeciesLM Fungi nucleotide reconstructions (scaled by information content) and nucleotide dependency map for the SMT3 promoter (yeast). Transcription factor (TF) motifs and poly(dA:dT) are reconstructed with similar confidence whereas blocks appear only for TF motifs in the dependency map. Ground truth motifs from http://www.yeastepigenome.org/. **B)** Examples of dependency blocks from human promoters. From top to bottom: Znf652 motif in the LDLR promoter, Nfy motif in the promoter of MTO1 and a Spdef motif in the ARID1B promoter. Ground truth motifs from Hocomoco v12 ^31^. **C)** Top. Per nucleotide block-score for nucleotides in repeats and reported to be in a bound TF motif. Bottom. Per nucleotide information content of the DNA-LM reconstruction in repeats and reported to be in a bound TF motif. **D)** Receiver operating characteristic (ROC) curve comparing the ability of different metrics to classify whether a nucleotide is part of a bound TF motif or not. The dependency block score performs significantly better than using the LM nucleotide predictions and is as good as yeast expert position weight matrix (PWM) scanning. This is even though PWMs were derived from in-vitro and in-vivo binding assays and were used for the definition of the positive class, whereas the language model has never been exposed to binding data during training. **E)** Dependency map for an instance of the yeast Abf1 spaced motif, compared to the ground truth binding preference from YeTFaSCo ^32^.

To find these dependency blocks we computed the first quartile of all query-target dependencies among consecutive 6 nucleotides (Methods). This quantile-based block score is more robust than the average to isolated strong interactions, privileging dense blocks. To benchmark the ability of the block score to serve as a binding-site detector, we leveraged the near-complete availability of TF binding data in *S. cerevisiae* along with TF binding nucleotide level preferences (position weight matrix, PWMs, ^32^). We considered the 1-kb regions 5’ of start codons and defined PWM matches occurring within 10 bp of an experimental binding peak as binding sites for 68 TFs ^33^.

While reconstruction varied widely for binding site nucleotides and for repeat elements, the block score of binding site nucleotides was generally higher than for nucleotides in repeats (Fig 2C). Consistent with this observation, the block score discriminated binding site nucleotides significantly better than reconstruction (Fig 2D). As a comparison, phylogenetic conservation inferred from alignment of 7 *Saccharomyces* species had no discriminative power at this task (Fig 2D) ^25^.

Moreover, the block score discriminated binding site nucleotides as effectively as PWM scanning. This result is remarkable since the block score was obtained in a completely unsupervised fashion using masked language modeling on genomic sequence alone, whereas the PWMs were not only derived from experimental data but also used to define the positive class. The true extent of this capability of DNA LMs is only apparent when inspecting the dependency map and not from the reconstructions. Thus, this analysis demonstrates both the ability of DNA LMs to detect regulatory elements as well as the utility of the dependency maps.

We note that not all motifs appear as complete blocks. *S. cerevisiae* Abf1, for example, is represented as two spaced and interacting blocks, correctly reflecting the dimeric binding preferences of this factor (Fig 2E). Thus, even within motifs, the dependency maps can serve to visualize the underlying functional relationships.

### Off-diagonal blocks highlight sequence element interactions

Blocks in the dependency maps also occurred away from the diagonal, typically reflecting distal dependencies between sequence elements, as for example between the TATA box and initiator (INR) element of the promoter of GstO2 in *Drosophila melanogaster* (Fig. 3A), two sequence elements involved in transcription preinitiation complex assembly, or between the donor site, the branch point and the acceptor site of the intron of ATG44 in *S. cerevisiae* (Fig. 3B), which are the three major sequence determinants of splicing. The short length of yeast introns allowed us to perform a genome-wide assessment which showed that dependencies between donor and acceptor splice sites were higher than dependencies between donor and decoy acceptor-like sequences within the intron or background dependencies at matched distances (Fig. 3C). These results indicate that distal dependencies capture a range of functional relationships between sequence elements, including promoter and transcript architecture.

**Fig. 3.**
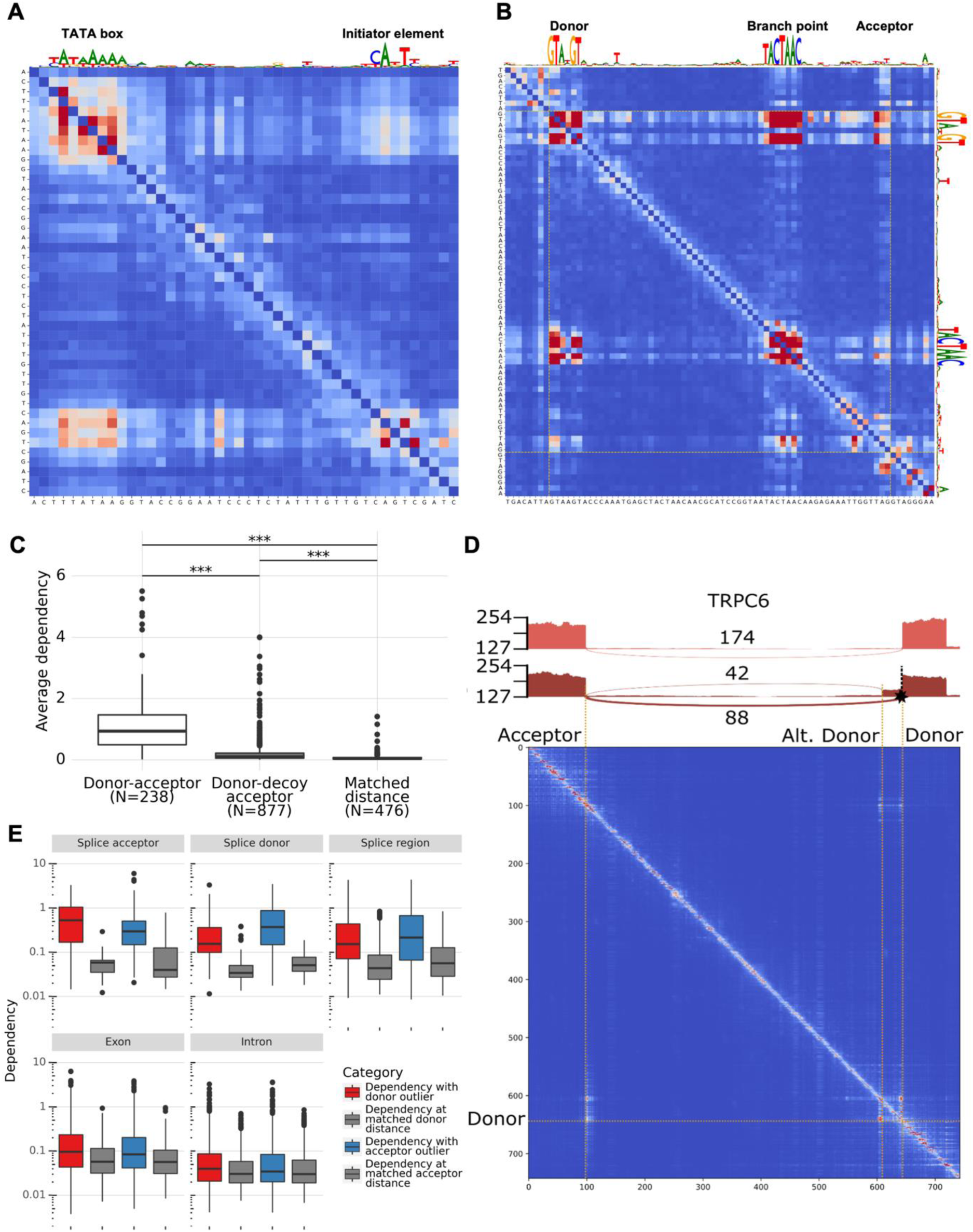
Off-diagonal blocks highlight sequence element interactions. **A)** Dependency map extracted from the SpeciesLM Metazoa in the promoter of the *Drosophila melanogaster* gene GstO2. On top is the reconstruction (scaled by the information content) from the LM highlighting the TATA box and initiator (INR) element motifs. High dependencies can be spotted at their intersection in the dependency map, reflecting their functional interaction. **B)** Dependency map for the intron of the yeast gene *ATG44* together with exonic flanking regions of 8 nucleotides. The top and right tracks correspond to the nucleotide reconstruction (scaled by the information content) which highlight the donor and branch point motifs. Off-diagonal dependencies can be spotted in the intersection between these motifs and the acceptor, indicating their interdependence. **C)** Average dependency between donor and acceptor nucleotides; donor and acceptor-like decoy nucleotides (AG dinucleotides within the intron not part of an annotated 3’ intron end); donor and random nucleotide pairs matching donor-acceptor distances. **D)** Exon elongation variant in a human individual on an intron of gene TRPC6. Top. Sashimi plots for an individual without the variant (upper track) and an individual with the variant (lower tracks) indicating differential splicing (number of RNA-seq reads supporting each splice junction) resulting from a variant inducing exon elongation. Bottom. Dependency map obtained from SpliceBERT showing a dependency between the canonical and alternative donor, indicating that a substitution in the canonical donor site induces a change in predicted probability for the alternative donor position shown in the Sashimi plots. **E)** For each variant location with respect to the splice site each boxplot shows: the average dependency between a variant position and its reported outlier junction donor or acceptor and average dependencies for nucleotides at distances matching the spacing between the variant and the outlier donor or acceptor. All comparisons between dependencies for donor and acceptor and dependencies at matched distances were significant (one-sided Wilcoxon rank-sum test, all P-values < 10^-12^).

Going a step further, we asked whether the maps could also reflect changes in transcript structure due to interindividual variation. To this end, we leveraged aberrant splicing events associated with rare variants from 621 human individuals (GTEx, ^34^) and SpliceBERT, a language model trained on vertebrate RNA sequences ^14^. The gene TRPC6 provides an example where one individual harbors a rare genetic variant disrupting a canonical donor splice site. In this individual, a cryptic intronic donor splice site is used instead, resulting in an aberrant, shorter, intron (Fig. 3D). Consistent with the effect of this variant on transcript structure, the canonical donor strongly interacts in the dependency map with both ends of the aberrant intron, namely the canonical acceptor and the cryptic donor (Fig. 3D). Across all 1,811 rare-variant associated aberrant splicing events amenable to this analysis, dependencies between the variant position and the ends of the corresponding outlier intron were higher than between nucleotides at matched distances (Fig. 3E). These results held for either end of the outlier intron and all location categories of the variant (Fig. 3E). In summary, these observations show that dependency maps capture splicing rules and can be indicative of transcript structure alterations resulting from genetic variants.

### Nucleotide dependencies reveal RNA secondary and tertiary structure contacts

Besides blocks, we also frequently noticed anti-parallel diagonals, i.e. distal stretches of consecutive nucleotides that depend on each other one-to-one in reverse order as in the case of the four arms of the yeast arginine tRNA described above (Fig. 1D). Using a simple convolutional filter, we systematically called regions with anti-parallel elements across the genomes of different fungi (Methods, Fig. S4A). Dependencies in anti-parallel diagonals were typically consistent with Watson-Crick or wobble base pairing (Fig. S4B), indicative that they captured helical stems, which are RNA structural elements that play a major role in determining RNA folding. Moreover, anti-parallel diagonals with the strongest dependencies were found among highly structured RNAs such as tRNAs and rRNAs (Fig. S4C, Methods). Hence, these findings suggest that detecting antiparallel diagonals in nucleotide dependency maps could be instrumental in inferring RNA structures.

To evaluate the potential of dependency maps for capturing RNA structures more broadly, we used RiNALMo, a language model trained on 36 million non-coding RNA sequences from a wide variety of species ^36^. As simple contact prediction scores, we retained the largest of the two dependency map entries for each pair of nucleotides. Despite these scores not being informed by prior structural data, they were strongly predictive of secondary structure contacts, with areas under the ROC curve typically exceeding 0.9 for most RNA families (using the Archive ll database ^37^, Fig. 4A).

**Fig 4.**
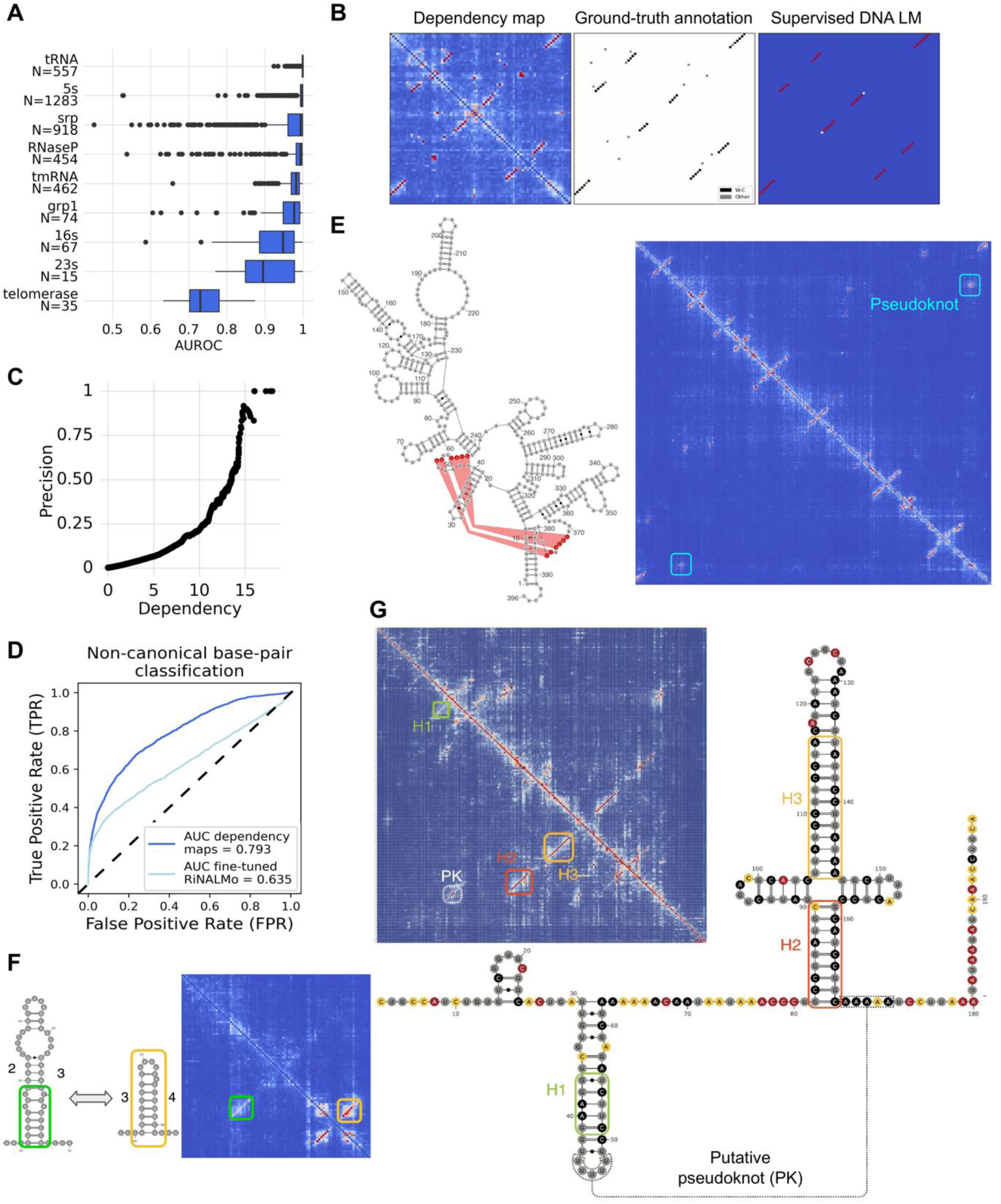
Dependency maps reveal known and novel RNA structures and highlight tertiary contacts. **A)** Area under the receiver operating characteristic curve (AUROC) for the classification of RNA structure contact pairs from the ArchiveII dataset spanning 9 different RNA families. **B)** *Archaeoglobus fulgidus* tRNA(Ile2) (PDB structure 3AMU) dependency map (left), ground-truth contacts (center), contacts predicted by the fine-tuned RiNALMo (right). **C)** Ratio of correctly retrieved contacts (Precision) not predicted by the supervised RiNALMo (predicted probability<0.5) for each dependency value threshold. **D)** ROC curve for the classification of non-canonical structure contacts across the CompaRNA dataset showing that dependency maps capture non-Watson-Crick and tertiary structure contacts which are lost on the supervised RiNALMo. **E)** Left. *Bacillus subtilis* RnaseP secondary structure highlighting pseudoknot contacts. Right. Corresponding dependency map showing anti- parallel dependencies belonging to RNA structure stems and the annotated pseudoknot contacts. The structure was taken from the RFAM database ^39^ with identification number RF00011. **F)** Tryptophan operon leader dependency map together with annotation and representation of the secondary structure stems belonging to sequence domains 2, 3 and 4. **G)** Left. Dependency map computed with RiNALMo of a region including 200 bp upstream of the gene *FkpB*. Right. DMS-MaPseq derived secondary structure together with reactivities per nucleotide. The DMS-MaPseq data are consistent with the dependency map. The main structural features are highlighted by boxes. Each stem-loop is identified starting with “H”, and the pseudoknot with “PK”. This structure was undescribed so far.

Originally, the authors of RiNALMo trained this language model as an intermediate step to train a supervised model specialized for predicting RNA secondary structures. Secondary structures are simplified planar representations of the topology of a single possible 3D folding of an RNA sequence. While informative, secondary structures miss important contacts occurring in the 3D fold. We noticed that some of the apparent false positive predictions of our dependency-map approach corresponded to tertiary structure contacts which were absent from the predictions of the supervised RiNALMo model. For instance, in the case of the *Archaeoglobus fulgidus* isoleucine tRNA, the dependency maps showed 6 base pairs with dependencies as strong as secondary-structure contact dependencies (dependency>6). These base pairs included 6 out of the 8 known contacts found in the tertiary structure that do not involve Watson-crick or wobble interactions, and therefore, are absent from the secondary structure. RiNALMo’s supervised model, whose task is to predict secondary structures, correctly captures the secondary structure contacts but not the remaining contacts found in the tertiary structure (Fig. 4B). This finding is interesting because the secondary structure of RNA, on its own, usually does not provide enough information to determine its 3D structure. Additional tertiary interactions offer useful spatial constraints that help in inferring the fold.

To systematically evaluate the added value of dependency maps to capture tertiary structure contacts we analyzed the database CompaRNA ^38^, which is based on available RNA structures from PDB. We found that 50% of the pairs with a dependency score larger than 13.5 and not predicted to be in secondary structure by Rinalmo’s supervised model were annotated as contacts (Fig 4C). CompaRNA annotates whether a nucleotide pair belongs to canonical or non-canonical base-pairing, with canonical base-pairing being cis Watson-Crick or wobble (G-U), which are the only contacts present in secondary structures. Across the entire database, non-canonical base pairs were well captured by the dependency maps (Fig. 4D, AUC 0.8). In contrast, this information was largely lost by the supervised model RINALMo trained on top of the language model (Fig 4D, AUC = 0.64, P < 10^-4^, permutation test). These results indicate that dependency maps can become a helpful tool for RNA structure inference by providing candidate contacts not captured by secondary structure contact predictors.

These findings prompted us to investigate further the potential of dependency maps in addressing the major challenges of secondary structure prediction. Among these, the prediction of pseudoknots still constitutes a largely unsolved problem in RNA computational biology. Pseudoknots are important non-secondary structure elements forming when base-pairs are not nested, such as when bases in a loop pair with a single-stranded region elsewhere on the RNA. We observed high dependencies between bases of documented contacts implied by pseudoknots. One striking example is shown in the 396 nt-long RNaseP RNA (Fig 4E, and Fig S4D for another example in a riboswitch), in which not only the stems but also the pseudoknot are reflected with strong antiparallel diagonals. These contacts were supported by co-varying bases in a sequence alignment, indicating that the structure has been conserved since a common ancestor sequence in present-time species.

An RNA’s secondary structure represents the topology of a single conformation. However, an RNA sequence can adopt alternative RNA folds to exert its functions. We found that dependency maps can capture alternative structures. For instance, the dependency maps of the tryptophan leader sequence in the bacterium *E. coli* (Fig. 4F), a structured region for which tryptophan abundance regulates the switch between terminator and antiterminator conformations ^40^, captures the two alternative folds, with the domain 3 being involved in anti- parallel diagonals with both domain 2 and domain 4 ^40^.

To assess the capacity of dependency maps to derive novel predictions, we performed in-cell chemical probing of *E. coli* with dimethyl sulfate followed by high-throughput mutational profiling analysis (DMS-MaPseq), a transcriptome-wide assay probing adenines and cytosines not engaged in Watson–Crick base-pairing ^41^. *E. coli* is well-suited to perform such studies due to the functional importance and prevalence of structured RNAs in bacteria and because sufficient sequencing depth can be obtained to have sensitive transcriptome-wide probing. We computed dependency maps for all non-coding regions upstream of the start codon spanning 500 nucleotides, as they are known to harbor different structures with roles in translation and transcription regulation ^42^. We selected dependency maps indicating the presence of at least 2 stem loops and not belonging to an annotated structure, revealing 4 previously unreported secondary structures corroborated by experimental data from DMS-MaPseq and validated by covariation analysis (Fig. 4G and Fig. S4G). Notably, as covariation analysis typically requires a high-quality sequence alignment and, optionally, a predicted RNA structure, the ability of nucleotide dependencies to capture – in an alignment-free and unsupervised fashion – functionally-relevant RNA structural contacts, underscores their predictive power.

Altogether, these results show that dependency-map analysis can overcome the typical challenges associated with RNA structure prediction, capturing both secondary and tertiary structure contacts, pseudoknots, and alternative structures of functionally-relevant RNAs.

### DNA LMs capture forward and inverted duplications without memorization

Analogous to antiparallel diagonals, we also frequently observed parallel diagonals in the dependency maps, as seen in the promoter of YNR064C in *S. cerevisiae* (Fig 5A). In this example, the sequence is a tandem repeat of two adjacent identical sequences of length 16. The parallel diagonal reflects that in the context of the tandem repeat, it is the identity of the n-th nucleotide that is predictive of the n-th nucleotide in the other repeat. This parallel diagonal pattern within the tandem repeat contrasts with the previously mentioned block pattern within regulatory sequence motifs (see TATA box, Fig. 5A) for which all nucleotides depend on each other.

**Fig. 5.**
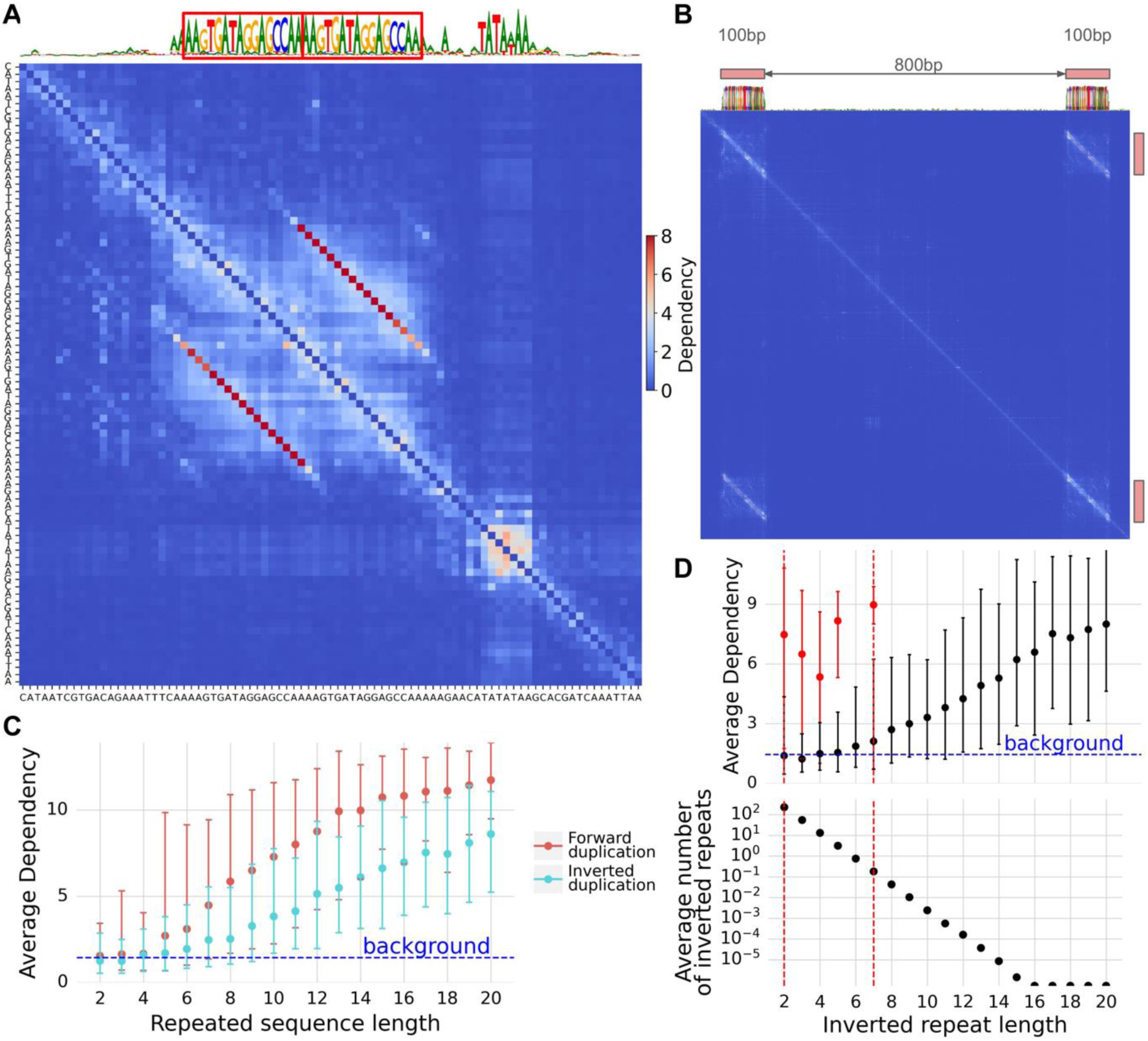
DNA LMs capture forward and inverted duplications without memorization. **A)** Dependency map from the SpeciesLM Fungi in the promoter of YNR064C which contains a TATA box motif and duplicated sequences highlighted as red boxes on top of the reconstruction. While the TATA box appears as a block-like dependency pattern, the repeat shows a parallel dependency linking each duplicated nucleotide. **B)** Dependency map and nucleotide reconstruction for a 1kb random sequence containing an inserted artificially generated random duplicated sequence of 100b. Despite the repeated sequences being spaced 800 bp apart, the DNA LM highlights the parallel dependency linking each nucleotide. **C)** Average dependency between artificially inserted repeat elements against their length for forward and inverted duplicates together (error bars: 95% confidence interval computed across 100 samples). **D)** Top. Average dependency against inverted repeat length for tRNA length sequences (Black colored dots). The red colored dots indicate the average dependency within anti- parallel dependencies in tRNA stems. Error bars indicate 95% confidence intervals. Bottom. Average number of inverted repeats expected to get by chance for each repeat length.

To study general properties of parallel diagonals, we first systematically scored dependencies for exhibiting a parallel diagonal pattern across the *S. cerevisiae* genome (Methods). The parallel diagonals with the strongest average dependencies were enriched for duplicated sequences (Fig. S5A). These observations suggested that the DNA language model did not memorize the sequence of the individual repeated elements but modeled the duplication itself. To test this hypothesis, we generated artificial sequences that include repeats of different lengths and spacing. As shown in Fig. 5B for two repeat lengths of 100 nucleotides spaced as far as 800 nucleotides from each other, the DNA LM reconstructed the randomly generated repeated sequences correctly with high confidence and showed the one-to-one nucleotide dependencies as parallel diagonals. Since the randomly generated sequences were not part of the natural training sequences, this observation shows that the DNA LM has learned the duplication operation. The capacity for identifying duplications strengthened with repeated element length, whereby the average dependencies within parallel diagonals longer than 7 nucleotides was significantly higher than background dependencies (Fig 5C).

Similarly, the DNA LM captured duplicated sequences in the reverse complement orientation (Fig 5C, using antiparallel dependencies). Since, as shown above, antiparallel diagonals reflect stems in RNA structures, we asked whether the DNA LM captured RNA stems as specific cases of reverse complement duplications. We tested this hypothesis by focusing on all 70 unique tRNA sequences of *S. cerevisiae.* For each tRNA, we generated 100 random sequences of matched length and nucleotide composition in which we inserted at non-overlapping locations an arbitrary sequence and its reverse complement (Methods). As for the same-strand duplications, the DNA LM captured the reverse-complement relationships with dependencies increasing with repeated element length (Fig 5D). However, the *S. cerevisiae* tRNA arms contain stems with short stretches of base-pairing spanning between 2 and 7 nucleotides. For these short lengths, the average dependencies on simulated reverse-complement sequences do not exceed twice the background dependencies and are substantially weaker than the average dependencies observed for the endogenous tRNA sequences (Fig 5D). This shows that the DNA LM does not rely only on reverse complementarity alone to link bases in contact and must leverage the broader context of the tRNA sequence. This property is important for functional contact predictions because randomly occurring short reverse complement sequences are pervasive (Fig 5D) and could lead to an overwhelming number of false positives.

### Dependency strength depends on genomic distance

We next investigated global properties on the distribution of dependencies, independently of specific patterns. To this end, we focused on *S. cerevisiae* as a model system. Nucleotide dependencies followed a power-law relationship with respect to distance to the query nucleotide, decaying by about 78% per 10-fold distance increase (Fig. 6A). We did not find substantial variations in the decay rate across various types of genomic regions (Fig. 6B). However, dependency maps in mitochondrial DNA showed a higher scaling constant than dependency maps from nuclear DNA regions, reflecting that dependencies were in general 1.64x stronger in the mitochondrial than in the nuclear genome (Fig 6B). Browsing dependency maps of mitochondria revealed regions of very rich and intricate dependencies whose biological interpretation needs further investigations (Fig. 6C for a representative example). Investigating deviations to the general power-law trend revealed higher dependencies at 3-nucleotide spacing, perhaps as a consequence of the high content of coding sequences in yeast. Nucleosome positioning also appeared to influence dependency distributions, with stronger dependencies than expected by the power law at distances corresponding to nucleosome position periodicity on both investigated yeast species *S. cerevisiae* (164 bp) and *S. pombe* (152 bp, fig 6D) ^43^. Altogether, these analyses showcase that nucleotide dependency maps offer a new avenue to study general constraints on genomic sequences.

**Fig 6.**
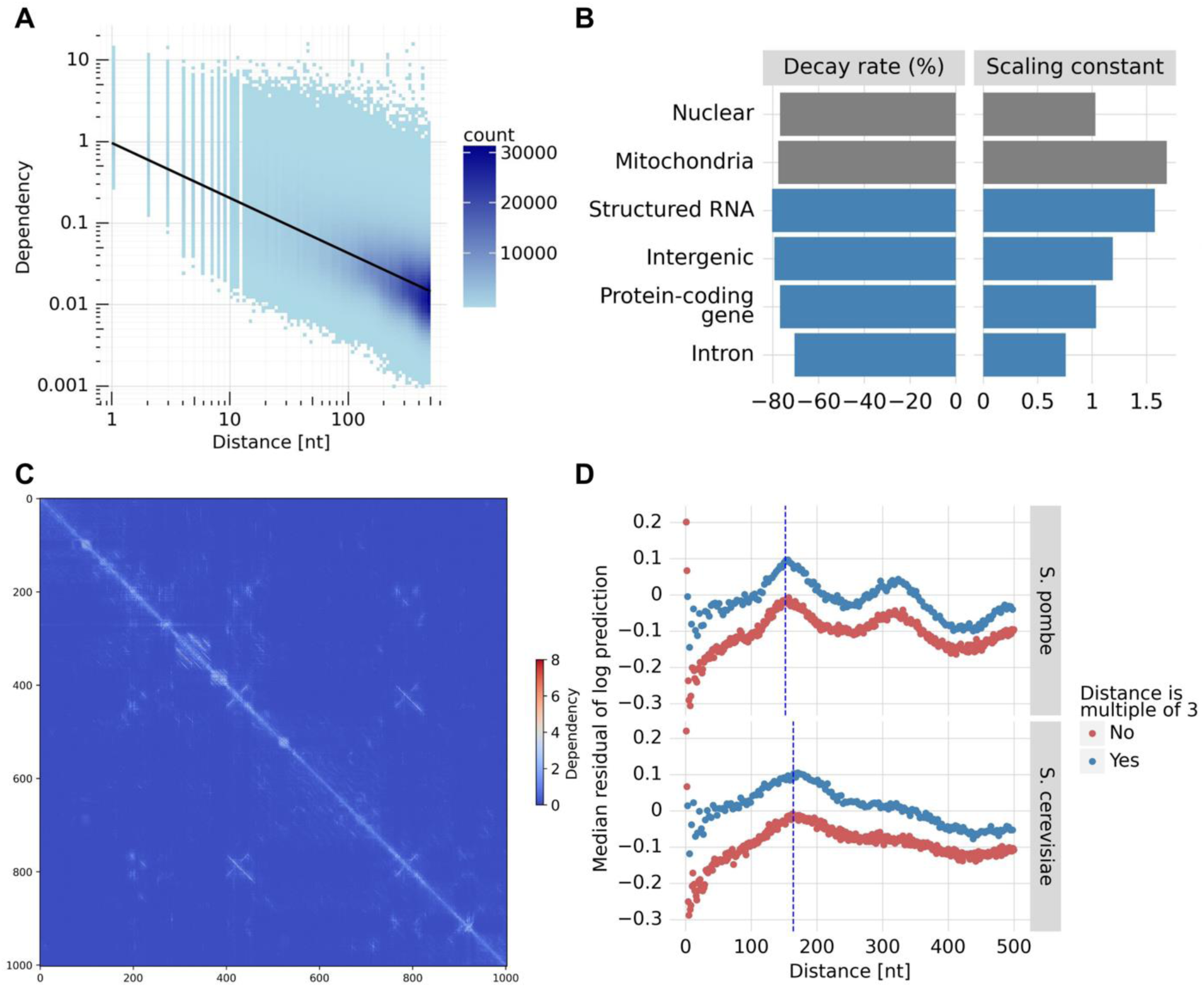
Dependencies relate to distance in a species and region-specific way and reveal periodicities intrinsic to the genome. **A)** Nucleotide dependencies computed from the SpeciesLM Fungi against distance to the query nucleotide taken from dependency maps across the yeast genome. Linear regression fit (black line) marks the power-law relationship between dependency and distance. **B)** Power-law decay rate in percentage decrease in dependency per 10-fold distance increase (left) and scaling constant (right) for different genomic regions (see Methods for exact category definitions). Example of a dependency map for the mitochondrial region 1kb 5’ of gene *Q0017*. **D)** Median of the residuals of the fitted power law against distance to the query nucleotide for yeasts *S. pombe* and *S. cerevisae.* The colors highlight the difference in dependency between targets distanced from the query by multiples of 3 nucleotides. The dashed blue lines show the nucleosome periodicity values reported for *S. cerevisiae* (164bp) and *S. pombe* (152bp) ^43^ and coincide with the highest deviations from the fitted power law, indicating that dependencies reveal and are constrained by nucleosome periodicity.

### Dependency maps uncover shortcomings in DNA LM model designs and training data selection

Current DNA LMs differ both in terms of model architecture and in terms of the sequence data they were trained on. As of writing, there is no consensus about the advantages and disadvantages of these different approaches. Such comparisons are difficult to perform since DNA LMs are extremely large and complex models, often employed as intermediate black boxes for training further models on top. We set out to use nucleotide dependencies, which can be computed for any DNA LM, as a general tool for visualizing and getting insights into existing DNA LMs.

Human tRNAs are suitable loci for performing such comparative diagnoses because several models have been trained on human genomes only and because tRNAs entail well-established and highly conserved distal functional dependencies. We observed that some modeling choices introduce artifacts in the dependency maps. For example, models belonging to the Nucleotide Transformer family ^9^ do not reconstruct at the single base level but instead predict non-overlapping spans of 6 nucleotides. This produces artificial dependency blocks along the diagonal, which do not represent motif instances but arise because nucleotides of the same span are generally more dependent (Fig 7A). Nevertheless, these models are capable of learning dependencies at the single base-level, such as evident for some tRNA stem contacts in the human tRNA-Arg-TCT-4-1 (Fig. 7A).

**Fig. 7.**
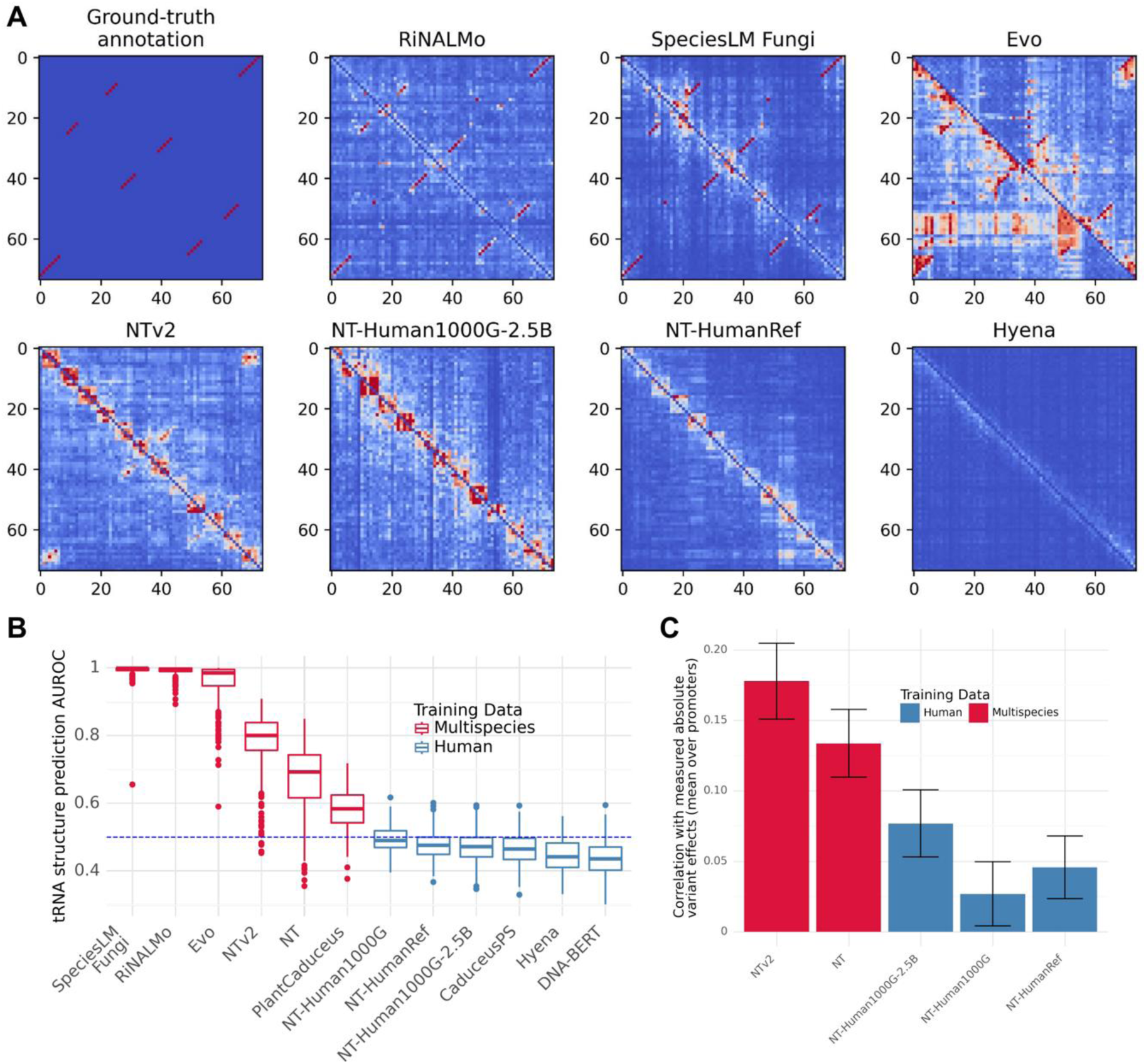
Dependency maps to compare DNA LMs and diagnose their shortcomings. **A)** The top left map shows the ground truth tRNA secondary structure contacts taken from GtRNAdb (tRNA-Arg-TCT- 4-1). Red indicates contact while blue indicates no contact. The remaining maps show the dependency maps for different DNA language models revealing modeling artifacts and performance differences. For instance, Nucleotide Transformer v2 ^9^ captures only a few of the structural interactions. The 6-by-6 blocks around the diagonal reveal an artifact of NTv2 non-overlapping 6-mer tokenization. Evo ^8^ shows artifacts when encountering the start of genomic elements (see upper left corner and lower-right corner). Computing the maximum across the upper and lower diagonal can mitigate the artifact. **B)** Comparing the AUROC achieved when dependency maps, as computed using different models, are used to predict secondary structure contacts of human tRNAs without fine-tuning. Models differ in terms of architecture and training data. Multispecies models strongly outperform those trained only on the human genome, even if the multispecies models have never seen human (or even metazoan) DNA. **C)** Comparing the correlation of the variant influence score, calculated using different Nucleotide Transformer models, with the measured absolute log fold change variant effect (compare Fig 1C). Multispecies models perform significantly better than those trained only on human sequences, even for models of the 1000G type which were exclusively trained across 3,202 diverse human genomes.

Equally, some models do not consider bidirectional context when making their predictions, but instead are designed to predict the next nucleotide given only its 5’ context. Evo is an example of such a so-called autoregressive model ^8^. This creates an artifact at the beginning of genomic elements such as the tRNA, which likely arises because the model cannot deduce the element until it has seen sufficiently many tokens inside of it (Fig 7A). At the cost of doubling the runtime, however, this problem can be mitigated by running the model both on the forward and reverse strand and taking the maximum dependency within a pair of nucleotides. Nevertheless, more appropriate measures of nucleotide dependencies for autoregressive models need to be developed such that the full right and left sequence context is considered.

Much starker differences are observed when comparing models trained on different types of sequences. Specifically, we found that models trained only on the human genome, regardless of architecture, parameter count, or whether within-species variation was included, did not learn the human tRNA structure to any meaningful degree (Fig 7B). By contrast, models trained on multiple species succeeded in at least learning aspects of human tRNA structure, regardless of architecture and whether the training data included any human genomes. To further investigate this point, we evaluated the performance of DNA LMs from the Nucleotide Transformer family, which all exhibit very similar architectures, on the human promoter saturation mutagenesis assay ^24^ (Fig 7C) and ClinVar ^23^ (SFig 7A). We again found that the multispecies versions of these models performed significantly better than the human-only ones, including models trained on thousands of different human genomes (Fig 7C). We conclude that infrequent genomic elements, even if they are highly conserved, generally require a multispecies approach to be learned.

## Discussion

In conclusion, we introduced nucleotide dependencies which quantify how nucleotide substitutions at one genomic position affect the likelihood of nucleotides at another position. This new metric, which can now be defined and computed thanks to the advent of DNA language models, appears as a general and effective approach to identifying functionally related nucleotides. Nucleotide dependency maps reveal functional elements across a wide range of biological processes including transcriptional, post-transcriptional regulatory elements, their interactions, and RNA folding. Therefore, this new metric has implications across multiple areas of computational and genome biology.

In the context of DNA LMs trained on multiple species, dependency relates to sequence conservation, a major indicator of functional importance leveraging purifying selection among homologous sequences, i.e. descending from a common ancestor sequence. In practice, sequence alignment is first used to identify homologous sequences; conservation is then estimated from the aligned nucleotide frequencies adjusted for phylogenetic drift and mutational biases. This approach limits the definition of conservation to alignable sequences. In contrast, DNA language models can more flexibly borrow information across sequences with similar contexts, allowing them to capture recurrent patterns such as transcription factor binding site motifs and their functional arrangements that can have arisen independently on non-homologous sequences. So far, alternative metrics proposed to substitute alignment-based conservation using DNA language models leverage reconstruction probability ^5,6^, based on the concept that unlikely sequences are more deleterious. We showed that the influence of a nucleotide’s identity on predicting others is a more effective indicator of deleteriousness and could outperform alignment-based conservation. However, we note that neither genetic drift nor mutational biases are controlled for in the dependency scores and may undermine their performance. Future research at the intersection of DNA language modeling and population genetics is needed to address these issues.

We have shown that dependency maps provide a promising novel entry point to unravel the regulatory code. Regulatory elements, such as transcription factor binding sites, manifest as dense blocks in dependency maps. We show in yeast that applying a simple image processing technique on dependency maps identified these sites with an accuracy comparable to models that required experimental binding data to be trained. This finding is important for unraveling regulatory elements for which experimental binding data are scarce, notably for post-transcriptional regulation and non-model species. Future work could address the limitations of the initial approach we proposed by allowing various sizes to the blocks or explicitly modeling the motifs underlying the blocks by exploiting the full base-level dependencies for each pair of positions. Beyond identification of regulatory elements, we have shown nucleotide dependencies highlight functional interactions between sequence elements in splicing and promoters. Further work could leverage nucleotide dependencies to derive functional relationships between regulatory elements to understand the sequence context in which they operate.

Dependency maps reflect bases in contacts in RNA folds remarkably well, a significant finding given the limited ground truth data in RNA structural biology. Our entirely unsupervised approach notably overcomes limitations of secondary structure inference yielding information on both canonical and non-canonical contacts, pseudoknots, and alternative folding. We validated several novel RNA structural predictions in *E. coli* experimentally. Analyzing nucleotide dependencies within RNA structure sequences is related to covariation analysis which identifies compensating substitutions between pairs of positions on an alignment as evidence for evolutionary conserved contacts. The dependency map approach alleviates the need for alignments, which are rarely unique and for which ambiguities, even by a single-nucleotide shift, affect the covariation statistics. However, it relies on the DNA language model to have been trained on enough homologous sequences of the RNA of interest to have captured these evolutionary footprints. In this respect, future work could investigate the influence on the choice of species, sequences, and model design.

Although DNA language models have hundreds of millions or billions of parameters their evaluations are often based on high-level aggregate statistics, such as area under the roc curve and R², assessing the performance on downstream tasks of further models that build on them. These evaluations conflate the contributions of DNA LMs as foundational models with those of the downstream supervised models and provide narrow, unidimensional assessments. Nucleotide dependencies offer a richer approach for visualizing the functional relationships uncovered by DNA LMs and enable benchmarking the DNA LMs themselves. We revealed critical limitations in current model architectures and single-species training practices, paving the way for more effective and generalizable DNA language models.

Across scientific fields, visualization tools allow researchers to make new observations and novel hypotheses. A non-quantifiable contribution of dependency maps, but perhaps not the least, might be to allow visualizing selective constraints on sequence in a novel way.

## Methods

### SpeciesLM Training

For SpeciesLM Metazoa, we obtained metazoan genomes comprising 494 different species from the Ensembl 110 database ^44^. For each annotated protein-coding gene, we extracted 2,000 bases 5’ to the start codon and trained a species aware masked language model on this region. We followed the training and tokenization procedure outlined in Species-aware DNA LMs ^4^, but kept the batch size at 2304 at increased input sequence length, resulting in about twice as many tokens seen during training as SpeciesLM Fungi 5’. We used rotary positional encoding to inject positional information into the Transformer blocks.

For SpeciesLM Fungi, we deviated from the above recipe by tokenizing each base of the sequences from Karollus et al. ^4^ separately (single nucleotide, 1-mer tokenization) and using learned absolute positional encodings. To stabilize training, we increased dropout in the MLP layers of the transformer to 0.2 and set it to 0.1 for attention dropout.

Overall, we improved training efficiency by fusing biases of the linear layers, the MLP in the transformer and the optimizer using Nvidia Apex. We used FlashAttention2 ^45^ to train all models.

### Nucleotide Dependencies and variant influence score

We define the dependency between a variant nucleotide 𝑘_𝑎𝑙𝑡_ at position 𝑖 and a target position 𝑗 as

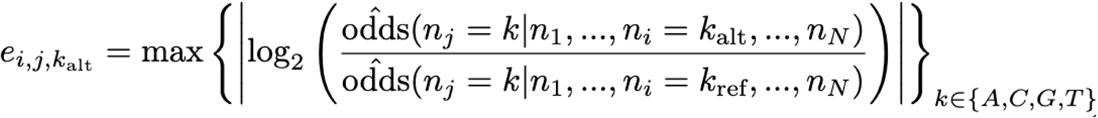

where 𝑘 is one of the 4 possible nucleotides A,C,G or T; 𝑛_𝑖_ and 𝑛_𝑗_ are the nucleotides at position 𝑖 and 𝑗 respectively; 𝑘_𝑟𝑒𝑓_ is the nucleotide in the reference, non-altered input sequence. The odds estimates are computed from the predictions of a DNA language model under consideration. For this computation, none of the nucleotides (including the target nucleotide) is masked.

The variant influence score 𝑒_𝑖,𝑘𝑎𝑙𝑡_, for a sequence of 𝑁 nucleotides is defined by averaging the dependencies on a variant nucleotide at position 𝑖 across all positions 𝑗 = 1, . . . , 𝑁 such that 𝑗 ≠ 𝑖 .

A nucleotide dependency 𝑒_𝑖,𝑗_ between a query position 𝑖 and a target position 𝑗 on a sequence of 𝑁 nucleotides is given by:

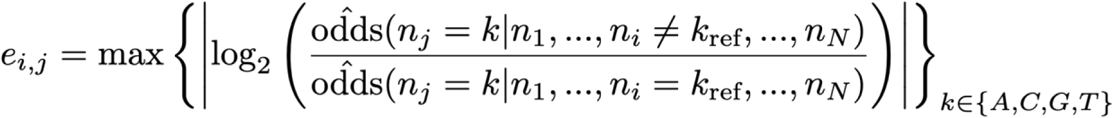

We compute dependencies for all 𝑖 , 𝑗 pairs such that 𝑖 ≠ 𝑗 , i.e. we do not consider self-dependencies.

In autoregressive models, a query variant cannot directly affect the prediction of a target position located 5‘ of the query. Thus, to get the lower triangular matrix of the dependency map, we run the model also on the reverse strand.

In the SpeciesLM Metazoa, which predicts nucleotides as overlapping 6-mers, the procedure needs to be adapted to yield one prediction per target nucleotide. This is achieved by first computing for each of the six 6-mers that overlap the target nucleotide of interest which probability it implies for this target nucleotide, as previously described ^4^. We then average these six probabilities to receive one probability.

For the Nucleotide Transformer models, which predict only non-overlapping 6-mers, we use a similar approach. Consider the case of predicting the probability of observing nucleotide 𝑛 at position 𝑖 of the sequence. In the tokenized sequence, this nucleotide has position 𝑝 in the k^th^ 6-mer where:

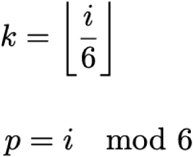

The model predicts a distribution over all 4^6^ possible 6-mers at position 𝑘. We first discard all predictions corresponding to 6-mers which contain a nucleotide that differs from the reference sequence at any location other than 𝑝 - which leaves only four 6-mers. We renormalize so that the predicted probability of these remaining 6-mers sums to one. We then record the (renormalized) probability of the 6-mer which has the desired nucleotide 𝑛 at position 𝑝.

Apart from extracting nucleotide level probabilities with the above-mentioned method, we have also experimented computing the probability for a nucleotide at position 𝑖 as the sum of all k- mers containing that nucleotide at that position. Evaluation of nucleotide dependencies within tRNAs revealed a worse performance with this method.

### Variant impact benchmarks

As our metric of variant impact, we used the variant influence score. This average is computed over the full receptive field of the model for the SpeciesLM. For Nucleotide Transformer models, we only average over the central 2 kb, so as to facilitate comparisons. Nevertheless, we provide the full sequence context this model has been trained for.

As a point of comparison, we also computed a variant effect score based on the DNA-LM reconstruction at the query variant. Specifically, this score is the log-ratio between the predicted probability of the variant nucleotide and the predicted probability of the reference nucleotide ^5,6^.

Lastly, we downloaded conservation scores (PhyloP and PhastCons) for Human and *S. cerevisiae* from UCSC ^25–28,46^. For human, these include the conservation scores based on the 100-way, 447-way and 470-way alignment.

### Promoter saturation mutagenesis

Promoter saturation mutagenesis data mapped to hg38 data was kindly provided by Vikram Agarwal. Following Avsec et al. ^30^, we excluded the FOXE1 promoter due to the low replicability of the measurements, leaving nine promoters and comprising 8,635 variants. Variants were then intersected with the human gene 5‘ regions (i.e. the regions 2 kb 5’ of annotated start codons). Then the variant influence score was calculated for each variant measured in the assay from the LM dependencies for these regions. The variant influence score was then correlated with the absolute value of the measured log2fold change in expression. This correlation was computed per promoter and then averaged over promoters.

To determine confidence intervals, we performed 100 bootstrap samples per promoter and recomputed the correlation for each bootstrap sample. The confidence interval was defined by adding/subtracting two standard deviations of the average correlation.

### eQTL variants

For human eQTL, we downloaded SUSIE ^18^ fine-mapped GTEx eQTL data from EBI. We then intersected this data with the human gene 5‘ regions. This procedure, by design, enriches for promoter eQTL. Similar to Avsec et al. ^30^, we considered every eQTL variant with a posterior inclusion probability higher than 0.9 as putative causal and we considered any eQTL variant with posterior inclusion probability lower than 0.01 as putative non-causal. We only considered putative noncausal eQTL intersecting regions which also include at least one causal eQTL.

This procedure gave 2,958 eQTL variants, of which 1631 were classified as putative causal. Then the influence score for each variant was computed from the nucleotide dependencies on these regions. We ranked variants according to the influence score. Confidence intervals were computed using bootstrapping as before.

For yeast eQTL, we downloaded the results of an MPRA study assessing candidate cis-eQTL variants ^20^. Following this study, we classify any eQTL variant with FDR < 0.05 in the MPRA assay as causal and we classify any eQTL with (unadjusted) P-value > 0.2 as non-causal. This yielded 3,056 eQTL variants of which 379 were classified as causal. These eQTL variants were then intersected with yeast gene 5‘ regions and influence scores were computed from the SpeciesLM Fungi dependency maps. Confidence intervals were computed using bootstrapping as before.

### Clinvar

We used ClinVar version 2023_07_17 ^23^, previously downloaded from https://ftp.ncbi.nlm.nih.gov/pub/clinvar/vcf_GRCh38/. We considered non-coding any variant in the categories ‘intron_variant’, ‘5_prime_UTR_variant’, ‘splice_acceptor_variant’, ‘splice_donor_variant’, ‘3_prime_UTR_variant’, ‘non_coding_transcript_variant’, ‘genic_upstream_transcript_variant’ and ‘genic_downstream_transcript_variant’. Following Cheng et al. ^47^, we considered as pathogenic any variant classified as pathogenic or likely pathogenic and as benign any variant classified as benign or likely benign. We excluded variants with fewer than one review stars. This resulted in 385,572 variants, of which 22,313 were classified as pathogenic.

As most ClinVar variants fall outside the 5‘ regions of genes, we chose not to intersect with these regions. Instead, we computed the dependency map centered on the variant of interest. Confidence intervals were computed using bootstrapping as before.

### Alternative dependency metrics

All benchmarks on alternative dependency metrics were performed on the SpeciesLM Fungi.

### Gradient-based

We computed the gradient of the prediction for each nucleotide at position 𝑖 with respect to each nucleotide at position 𝑗 yielding a 4x4 matrix. To achieve this, we first replaced the tokenization layer with a one-hot encoding and a linear layer, which maps the one-hot encoded nucleotides to their respective token embeddings. We then propagated gradients from each target nucleotide prediction to each one-hot encoded input nucleotide. As a metric of nucleotide dependency we then used the maximum absolute value across the 4x4 matrix of each 𝑖, 𝑗 position.

### Mask-based

Masked-based dependencies are computed as:

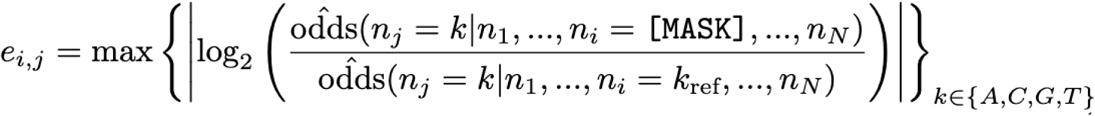

where [MASK] stands for the mask token, 𝑘 belongs to one of the 4 possible nucleotides A,C,G or T; 𝑛_𝑖_ and 𝑛_𝑗_ are the nucleotides at position 𝑖 and 𝑗 respectively; 𝑘_𝑟𝑒𝑓_ is the nucleotide in the reference, non-altered input sequence.

### *S. cerevisiae* tRNA structure benchmark

*S. cerevisiae* genome assembly version R64-1-1 and annotation version R64-1-1.53 were downloaded from EnsemblFungi ^44^. The *S. cerevisiae* tRNA secondary structures were downloaded from GtRNAdb ^48^. We considered only the tRNAs overlapping the 1-kb 5’ regions to any yeast start codon, yielding 172 tRNA sequences. Subsequently, dependency maps on tRNAs were processed by taking the maximum between 𝑒_𝑖,𝑗_ and 𝑒_𝑗,𝑖_ which symmetrizes the dependency map and achieves one unique score per pair of positions in the tRNA sequence. We then used this score to predict whether a pair belonged to a secondary structure contact.

### Assessment of donor-acceptor dependencies in *S. cerevisiae*

We extracted intron sequences by selecting the regions within annotated gene intervals that lie between exon annotations. This resulted in 380 sequences. We then retained only introns bounded by canonical splice site dinucleotides GT and AG, yielding 272 sequences. We then computed the average dependency between every donor and acceptor nucleotides within the intron as a measure of dependency between the donor and acceptor sites. We designed two negative sets for a given intron. For the negative set “Decoy acceptor” we compute the average dependency between donor nucleotides and each AG dinucleotide within the intron that does not include the acceptor site. For the negative set “Matched distance” we sampled four random dependencies between nucleotides that were as distant from each other as the donor was from the acceptor, without including the donor or acceptor themselves.

### TF motif mapping

We downloaded FIMO PWM scan results from http://www.yeastss.org ^49^ and Chip-Exo transcription factor binding peaks from http://www.yeastepigenome.org ^33^. We then extracted all Chip-Exo peaks for the available PWMs. We excluded PWM matches for which no Chip-Exo data for the corresponding factor was available. This procedure yielded data for 68 TFs. We annotated every nucleotide within 1 kb 5’ of a start codon as part of a binding TF motif if it is (1) part of a PWM match with P-value < 0.01 and (2) this PWM match is within 10 bases of a Chip-Exo peak of the corresponding transcription factor. We defined the positive class in this way to ensure that we capture nucleotides relevant for determining binding (i.e. motif) rather than all nucleotides close to a Chip-Exo peak regardless of their function in binding. This gave 92,117 binding nucleotides out of 6,538,427 overall. We designated a nucleotide as repeat if it was masked by RepeatMasker. We extracted this information from the soft-masked GTF provided by Ensembl ^44^.

### Dependencies in rare-variant-associated aberrant splicing

We computed dependency maps for all rare single nucleotide variants associated with splicing outliers in GTEx ^29^ as described previously ^34^. Since the input length of SpliceBert ^14^ is limited to 1,024 bp, the complete set of variant outlier pairs (N = 18,371) was filtered such that the variant and associated outlier junction were located within a 800 bp window (N = 1,811) and 100 bp of sequence was added from the maximum and minimum positions of the variant and outlier junction splice sites. For each variant location we extracted the average value of the dependency map at the intersection of variant and outlier donor dinucleotide or variant and outlier acceptor dinucleotide. This variant effect score was compared against a background score. This background score was computed as the mean over all dependencies that were as distant from each other as the variant was from the outlier donor (matched distance) or the outlier acceptor. The scores were filtered for a minimum distance of 5 bp between the variant and splicing dinucleotide to filter values near the diagonal corresponding to self interactions. Variant categories were annotated with the Ensembl Variant Effect Predictor (VEP) ^50^. For each variant, the most severe VEP annotation was considered. For the ‘Exon’ category, the following VEP categories were grouped together: synonymous_variant, missense_variant, stop_lost, stop_gained.

### Genome-wide search for parallel and anti-parallel dependencies

We scanned dependency maps for parallel and anti-parallel dependencies using 5x5 convolutional filters. We constructed the anti-parallel filter by populating the anti-diagonal of a zero-filled 5x5 matrix with ones, and for the parallel filter, by populating the diagonal with ones. We then centered each filter by subtracting the mean value from each position to ensure that a convolution on a uniform 5x5 region yields a result of zero. We applied these filters to dependency maps from SpeciesLM Fungi (both filters) and RiNALMo (anti-parallel filter only)4,36.

### Search for parallel and anti-parallel dependencies in Fungi using the SpeciesLM Fungi

For the SpeciesLM Fungi we have computed dependency maps spanning 1 kb 5’ of each annotated start codon on a set of representative fungi species including *Agaricus bisporus*, *Candida albicans*, *Debaryomyces hansenii*, *Kluyveromyces lactis*, *Neurospora crassa*, *Saccharomyces cerevisiae*, *Schizosaccharomyces pombe* and *Yarrowia lipolytica*. The genomes and annotation files for each species were downloaded from EnsemblFungi release 53 with accession numbers: GCA_000300555.1, GCA000182965v3, GCA_000006445.2, GCA000002515.1, GCA_000182925.2, GCA_003046715.1, GCA_000002945.2 and GCA_000002525.1 respectively.

All regions annotated as ’five_prime_utr’, ’three_prime_utr’, ’intron’, ’CDS’, ’pseudogene_with_CDS’ and other regions (ex. non-annotated introns) inside an annotated gene interval were categorized as Protein coding gene. All regions annotated as ‘tRNA’, ’tRNA_pseudogene’, ’rRNA’, ’snRNA’, ’ribozyme’, ’SRP_RNA’, ’snoRNA’, ’RNase_P_RNA’, ’RNase_MRP_RNA’ were categorized as Structured RNA. Finally, all regions annotated as ‘’transposable_element’, ’pseudogene’ as well as regions without any annotation were considered as intergenic.

### Search for anti-parallel dependencies and RNA structure in *E. coli* using RiNALMo

For RiNALMo we computed dependency maps for regions 100, 200 and 500 bp before each annotated start codon in *Escherichia coli* str. K-12 substr. MG1655 whose genome and annotation were downloaded from GenBank ^51^ with accession number U00096.3.

As candidates for novel RNA structure we first filtered positions whose convolution value is greater or equal to 25 to select only high value anti-parallel dependencies, resulting in a filtered convolved dependency map. Next, we counted the unique number of anti-diagonals potentially belonging to one stem by extracting the unique 𝑖 + 𝑗 non-zero positions supported by at least three non-zero values.

As candidates for novel structure we selected maps suggesting the existence of at least two potential stems.

### RNA secondary structure benchmarking

We downloaded the database of secondary structures Archive II ^37^, which includes 3,865 curated RNA structures across nine families (5S rRNA, SRP RNA, tRNA, tmRNA, RNase P RNA, Group I Intron, 16S rRNA, telomerase RNA, 23S rRNA). For each structure, we generated the dependency map with the pretrained RiNALMo and retained the largest of the two dependency map entries for each pair of nucleotides (maximum of 𝑖, 𝑗 and 𝑗, 𝑖 ). The area under the ROC curve was computed for each structure against the Archive II secondary structure annotations.

### Benchmarking of canonical and non-canonical RNA contacts

We downloaded the database of RNA structures CompaRNA ^38^, which is a compilation of RNA contacts based on 201 available RNA structures in the Protein Data Bank by RNAView ^52^. Contacts are classified either as "standard" or "extended". While the first includes only canonical AU, GC and wobble GU pairs in the cis Watson-Crick/Watson-Crick conformation ^53^, the latter calls all interacting bases regardless of their conformation, including non-canonical or tertiary contacts. Out of the 201 structures, 196 had a length below the maximum input length of RiNALMo (1,022 nt). For each structure, we generated the dependency map with the pretrained RiNALMo, and retained the largest of the two dependency map entries for each pair of nucleotides. Similarly, the same structures were also evaluated with the fine-tuned RiNALMo model version rinalmo_giga_ss_bprna_ft resulting in a predicted value per pair of nucleotides. To evaluate their performance on predicting non-canonical contacts, we excluded all canonical contacts and computed the area under the ROC curve of all remaining positions of all structures. Significance between ROC AUCs was determined by bootstrapping over 10,000 permutations.

### DMS-MaPseq analysis of E. coli cells

Escherichia coli TOP10 cells were grown in LB broth at 37°C with shaking until OD600 = 0.5, after which dimethyl sulfate (DMS, Sigma Aldrich, cat. D186309), prediluted 1:4 in ethanol, was added to a final concentration of 200 mM. Bacteria were incubated for 2 minutes at 37°C, and reaction was quenched by addition of 0.5 M final DTT. Bacteria were pelleted by centrifugation at 17,000g for 1 minute at 4°C, after which they were resuspended in Cell pellets were resuspended in 12.5 μl Resuspension Buffer [20 mM Tris-HCl pH 8.0; 80 mM NaCl; 10 mM EDTA pH 8.0], supplemented with 100 μg/mL final Lysozyme (cat. L6876, Merck) and 20 U SUPERase•In™ RNase Inhibitor (cat. A2696, ThermoFisher Scientific), by vortexing. After 1 minute, 12.5 μl Lysis Buffer [0.5% Tween-20; 0.4% Sodium deoxycholate; 2 M NaCl; 10 mM EDTA] were added, and samples were incubated at room temperature for 2 additional minutes. 1 mL TRIzol™ Reagent (cat. 15596018, ThermoFisher Scientific) was then added, and RNA extracted as per manufacturer instructions. rRNA depletion was performed on 1 μg total RNA using the RiboCop for Bacteria kit (cat. 126, Lexogen). DMS-MaPseq library preparation was performed as previously described ^41^. After sequencing, reads were aligned to the E. coli str. K-12 substr. MG1655 genome (GenBank: U00096.3), using the *rf-map* module of the RNA Framework ^54^ and Bowtie2 ^55^. Count of DMS-induced mutations and coverage and reactivity normalization were performed using the *rf-count*-*genome* and *rf-norm* modules of the RNA Framework. Experimentally-informed structure modeling was performed using the *rf-fold* module of the RNA Framework and ViennaRNA v2.5.1 ^56^.

### RNA structure covariation analysis

Covariation analysis was performed using the *cm-builder* pipeline (https://github.com/dincarnato/labtools) and a non-redundant database of 7,598 representative archaeal and bacterial genomes (and associated plasmids, when present) from RefSeq ^57^.

### Evaluation of artificial forward and inverted duplications

We generated random sequences of 100 nucleotides by sampling from regions 1 kb 5’ of the start codon in *S. cerevisiae* to ensure a representative GC content and shuffling the sequences to destroy potential functional elements. Additionally, we created 100 unique duplicated sequences, ranging from 2 to 20 nucleotides in length, by randomly sampling each nucleotide with equal probability. Each duplicated sequence was then inserted into a uniquely generated 100-nucleotide sequence at a random distance from each other, ensuring no overlaps occurred. We used the SpeciesLM Fungi to generate dependency maps for each sequence. We then computed average dependencies by taking the mean of the dependencies between nucleotides and their duplicates; this involved averaging across a parallel diagonal for forward duplications and an anti-parallel diagonal for inverted duplications.

For tRNA-sized sequences, we followed a similar method but generated each sequence by shuffling each unique tRNA sequence in *S. cerevisiae* once. We computed the average number of inverted duplications by averaging the occurrences of duplicated sequences of specific lengths across 10,000 shuffled versions of each tRNA sequence.

### Genome-wide analysis of dependencies distribution

Using the SpeciesLM Fungi we computed dependency maps across the genomes of *S. cerevisiae* and *S. pombe.* Since the SpeciesLM Fungi was pre-trained on sequences of 1,003 nucleotides including the start codon at the end, we discarded dependencies involving the 3 last nucleotides of each sequence yielding dependencies for 1,000 nucleotides. Genome-wide dependency maps of 1-kb span were obtained with a tiling approach. Along each chromosome, we computed 1-kb square dependency maps every 500 bp and averaged overlapping entries.

To ensure the same number of targets are computed before and after a specific query nucleotide, we considered dependencies involving nucleotides at most 500 positions away from each other. For each map we sampled 1,000 dependencies. Due to limitations in numerical precision we considered only dependencies larger than 0.001.

To compute the power law coefficients, a linear regression was fitted to predict the logarithm of the dependency from the logarithm of its corresponding distance in nucleotides. The scaling coefficient was then obtained by exponentiating the fitted intercept of the linear regression, and the decay rate was obtained directly from the fitted slope. The scaling coefficient and decay rate were computed for different regions in the genome: Nuclear - involving all dependencies belonging to nuclear DNA; Mitochondria - involving all dependencies within mitochondrial DNA; Structured RNA - belonging to the annotations ‘tRNA’, ’tRNA_pseudogene’, ’rRNA’, ’snRNA’, ’ribozyme’, ’SRP_RNA’, ’snoRNA’, ’RNase_P_RNA’ or ’RNase_MRP_RNA’; Protein-coding gene - belonging to the annotations ’five_prime_utr’, ’three_prime_utr’, ’CDS’ or ’pseudogene_with_CDS’; Intron - belonging to the regions inside an annotated gene interval but not to exons; and Intergenic - belonging to all regions annotated as ‘’transposable_element’, ’pseudogene’ as well as regions without any annotation.

### Model Comparison

All other models used were downloaded from Huggingface or from their publicly available repositories. Human tRNA sequences were downloaded from GtRNAdb ^48^. Exact duplicate sequences were removed, leaving 266 tRNAs.

## Data and code availability

Data and SpeciesLM model weights are available at https://zenodo.org/doi/10.5281/zenodo.12982536. Code is available at https://github.com/gagneurlab/dependencies_DNALM. Raw DMS-MaPseq data has been deposited to the Gene Expression Omnibus database (GEO), under accession GSE271937.

## Acknowledgements

P.T. is funded by the Munich Center for Machine Learning (MCML). N.W. is supported by the Helmholtz Association under the joint research school ‘Munich School for Data Science – MUDS’. D.I. was supported by the Dutch Research Council (NWO), NWO Open Competitie ENW – XS, project number OCENW.XS22.1.015 and by the European Research Council (ERC), European Union’s Horizon Europe research and innovation programme, grant agreement number 101124787, RNAStrEnD. X.H. was supported by an EMBO Postdoctoral Fellowship (ALTF 792-2022). J.G. was funded by the Deutsche Forschungsgemeinschaft (DFG, German Research Foundation) via the project NFDI 1/1 “GHGA - German Human Genome-Phenome Archive” (#441914366), by the German Bundesministerium für Bildung und Forschung (BMBF) through the ERA PerMed project PerMiM (01KU2016B) and through the Model Exchange for Regulatory Genomics project MERGE (031L0174A). J.G. was also funded via the EVUK programme (“Next-generation Al for Integrated Diagnostics”) of the Free State of Bavaria. J.G. and G.G. by the Deutsche Forschungsgemeinschaft (DFG, German Research Foundation) through the TRR267 (403584255, sub-project Z03). This study was supported by the Deutsche Forschungsgemeinschaft (DFG, German Research Foundation) via the IT Infrastructure for Computational Molecular Medicine (project #461264291). This study was funded by the European Research Council (ERC) (EPIC, Grant number: 101118521). Funded by the European Union. Views and opinions expressed are however those of the author(s) only and do not necessarily reflect those of the European Union or the European Research Council Executive Agency. Neither the European Union nor the granting authority can be held responsible for them.

