## Supplementary Material for "Nucleotide dependency analysis of DNA language models reveals genomic functional elements"

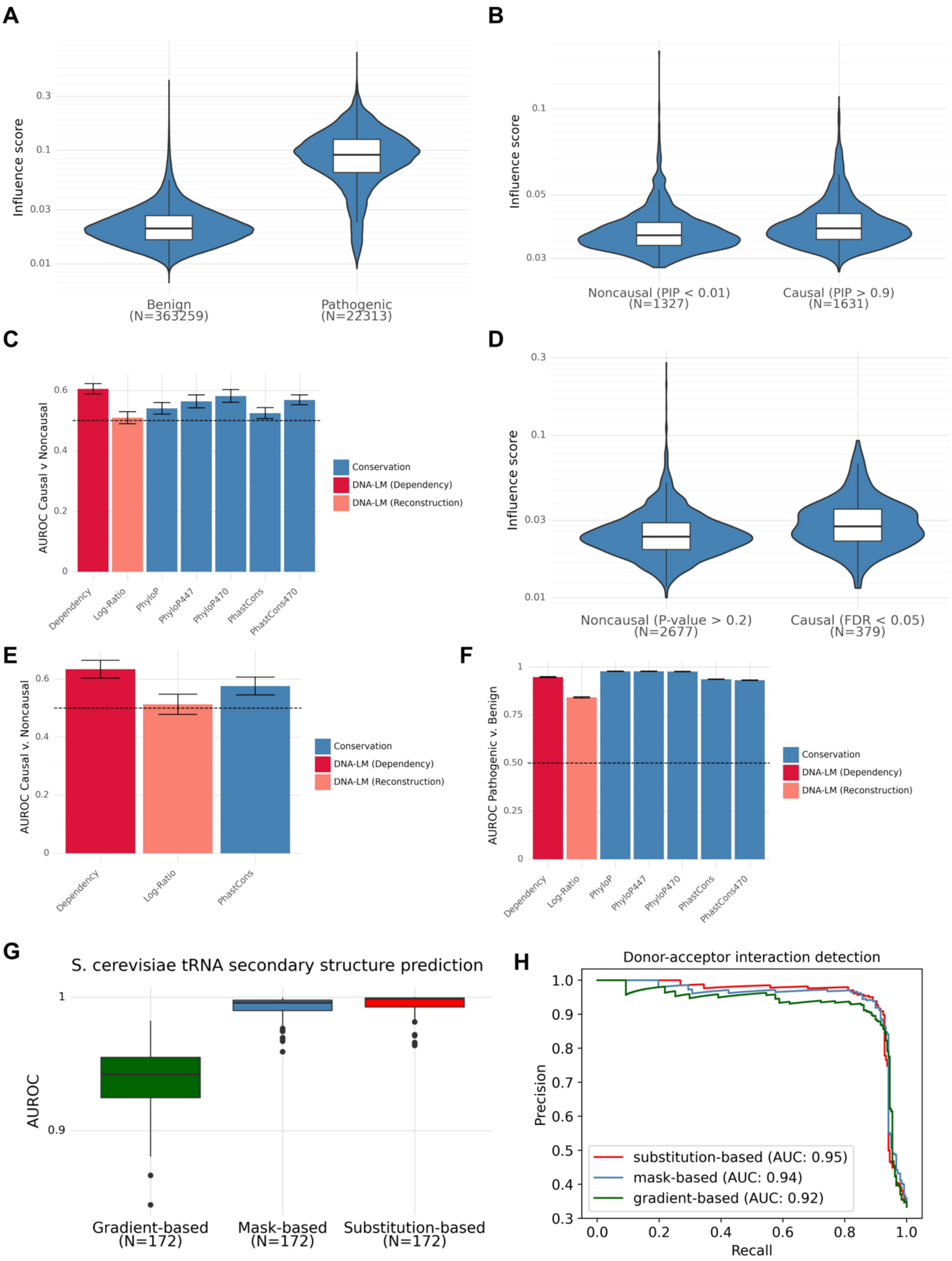


**Supplementary Figure 1. A)** Variant influence score against pathogenic and benign variants classified from ClinVar. P-value obtained from double-sided Wilcoxon rank-sum test <$10^{-6}$. **B)** Variant influence score for putative causal and putative non-causal variants as obtained from fine-mapped human eQTL [18,19]. P-value obtained from double-sided Wilcoxon rank-sum test <$10^{-6}$. **C)** Area under the receiver operating characteristic curve (AUROC) for the classification of putative causal versus putative non-causal variants from the Finemapped human eQTL comparing the variant influence score, DNA LM log ratio between predicted probability of the reference nucleotide and variant nucleotide, and alignment-based conservation scores from PhyloP and PhastCons. **D)** Variant influence score for putative causal and putative non-causal variants obtained from yeast eQTL [20]. P-value obtained from double-sided Wilcoxon rank-sum test $<10^{-6}$. **E)** Performance in AUROC for the classification of yeast putative causal vs putative non-causal eQTL variants. **F)** Performance (AUROC) for the classification of ClinVar variants into pathogenic or benign. **G)** Performance (AUROC) for the prediction of tRNA secondary structure contacts using different nucleotide dependency metrics: gradient-based, mask-based and substitution-based. **H)** Precision-recall curves for the prediction of splice site interactions using different nucleotide dependency metrics as before.

**A**

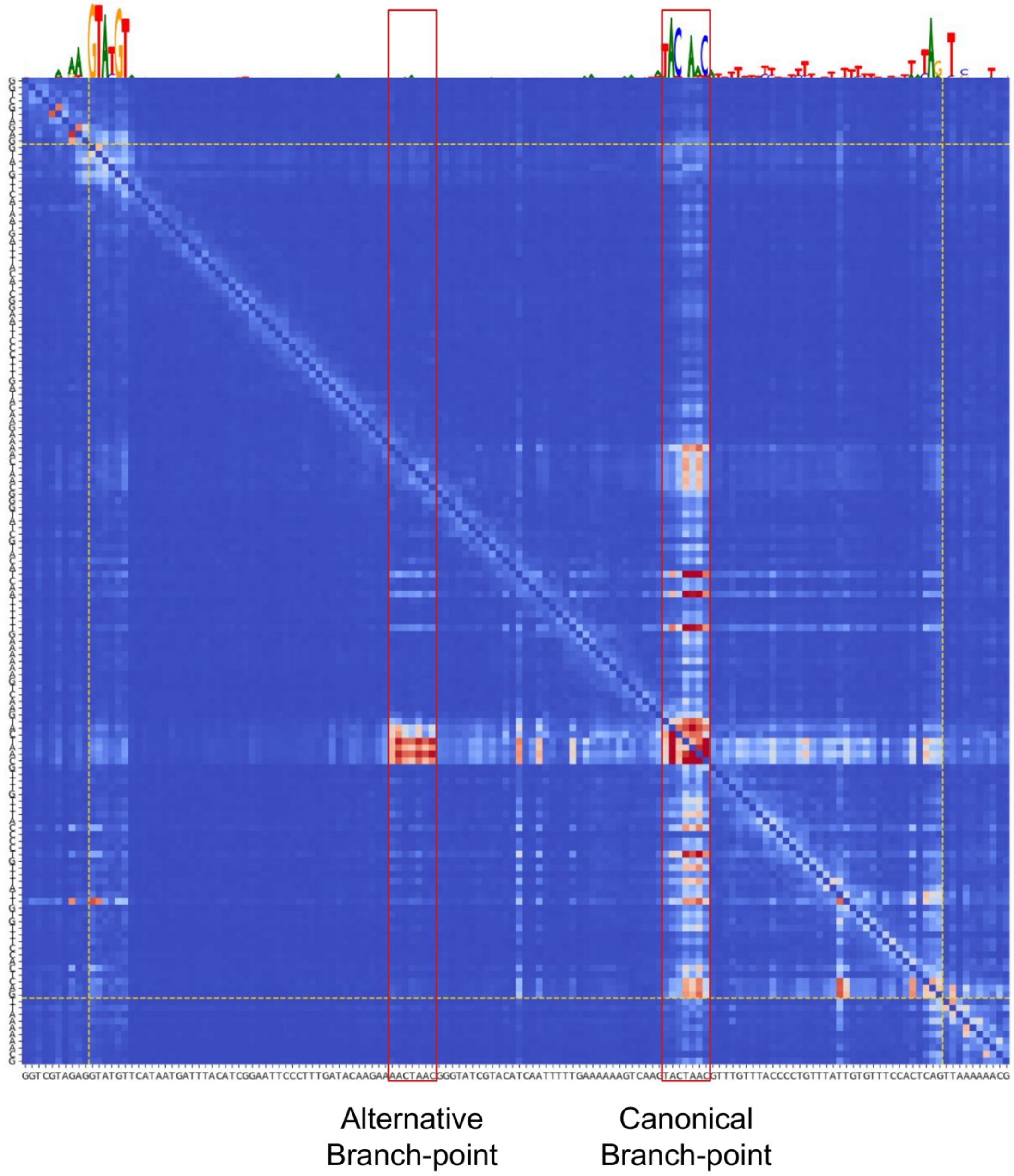


**Supplementary Figure 3. A)** Dependency map for an intron of the yeast gene *LSM2* not only highlighting the canonical donor, acceptor and branchpoint but also an alternative non-canonical branch point. While the canonical branch point appears as an on-diagonal block, another parallel off-diagonal block is visible, suggesting that if mutations altered the canonical branch point then compensatory mutations on the alternative branch points would be favored. The target nucleotides of this block belong to a branch-point-like sequence, indicating a role as an alternative branch-point which has also been previously found experimentally [35].

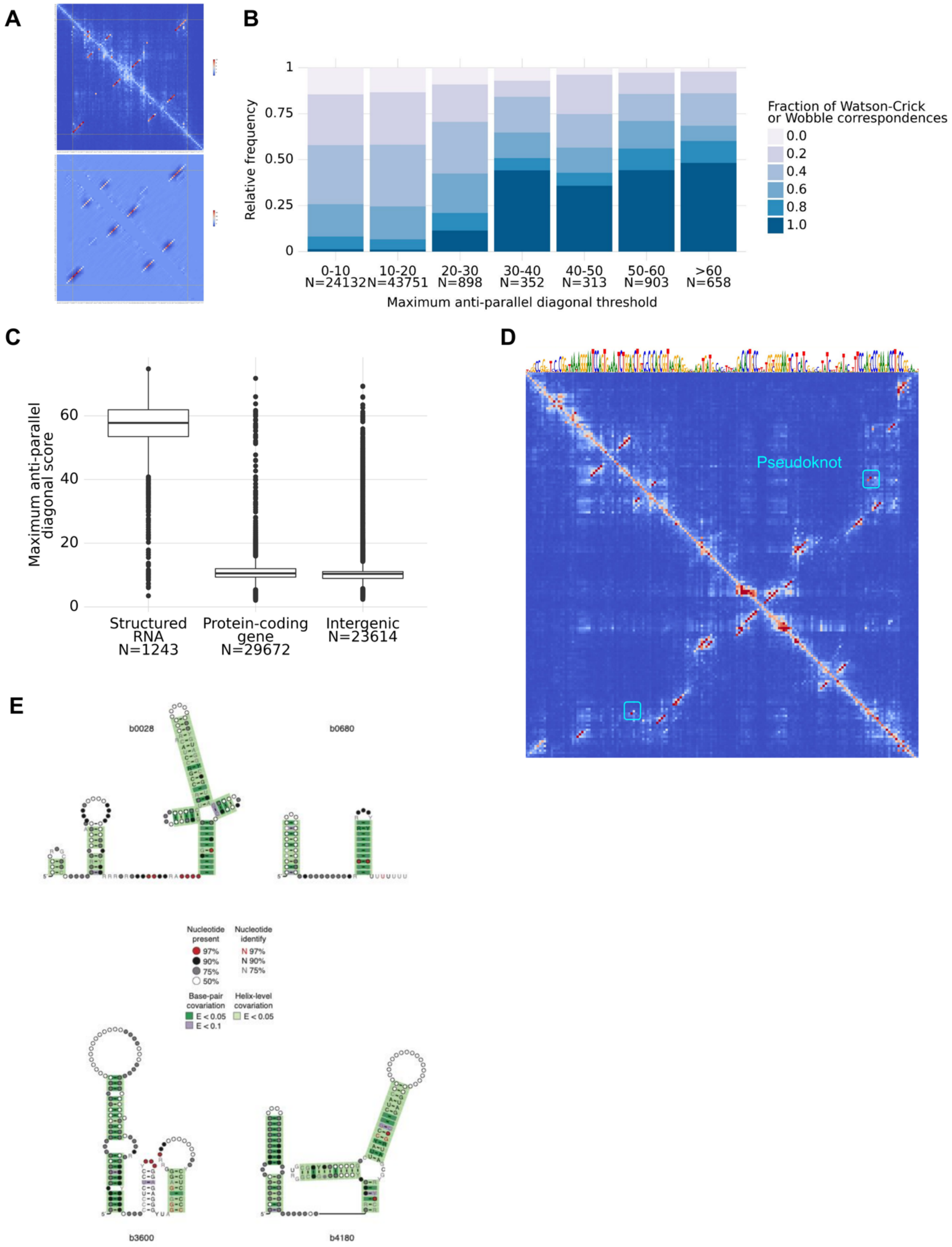


**Supplementary Figure 4. A)** Example of a convolution of a 5x5 anti-parallel diagonal filter on the dependency map of yeast tR(ACG)O tRNA. The dependency map is shown on top while the resulting convolution is shown in the bottom. The maximum convolution values highlight the anti-parallel dependencies within the tRNA. **B)** Fraction of nucleotides that show Watson-Crick or wobble correspondence within the 5 base-pair region defined by the convolution filter location with the maximum hit for different convolution values in a region of 1kb 5' of a start codon. **C)** Maximum anti-parallel dependencies within a 1kb region 5' of each start codon across fungi species and its location in the genome categorized in one of Structured RNA; Protein coding - spanning the whole interval of a protein-coding gene; and Intergenic - spanning mostly non annotated regions between genes. **D)** *E. coli* Cobalamin riboswitch dependency map together with the highlighted pseudoknot contacts and nucleotide reconstruction on top. **E)** Covariation analysis of four novel structures validated by DMS-MaPseq 5’ of genes FkpB (b0028), glnS (b0680), mtlD (b3600) and rlmB (b4180).

**A**

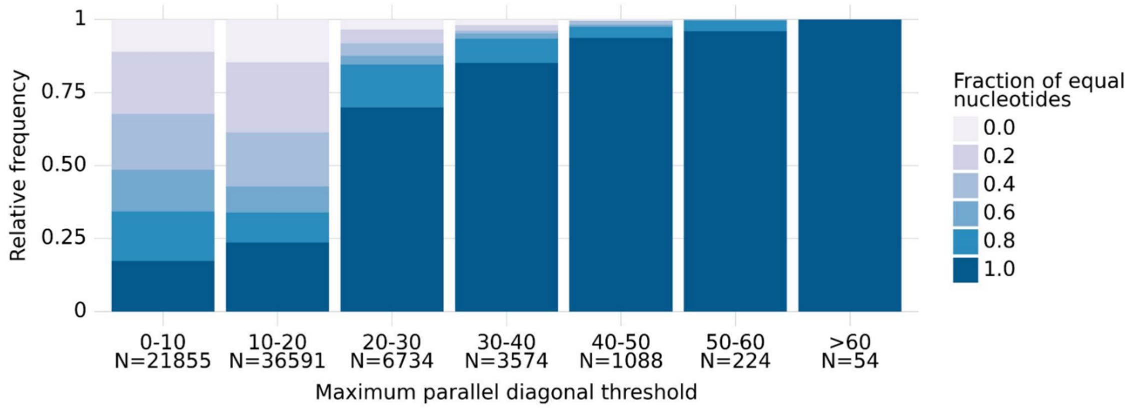


**Supplementary Figure 5. A)** Fraction of equal nucleotides within the 5 base-pair region defined by the parallel diagonal convolution filter location with the maximum hit for different convolution values. The strongest parallel diagonal dependencies belong to repeated sequences, indicating that repeats are highlighted genome-wide in parallel dependency patterns.

A

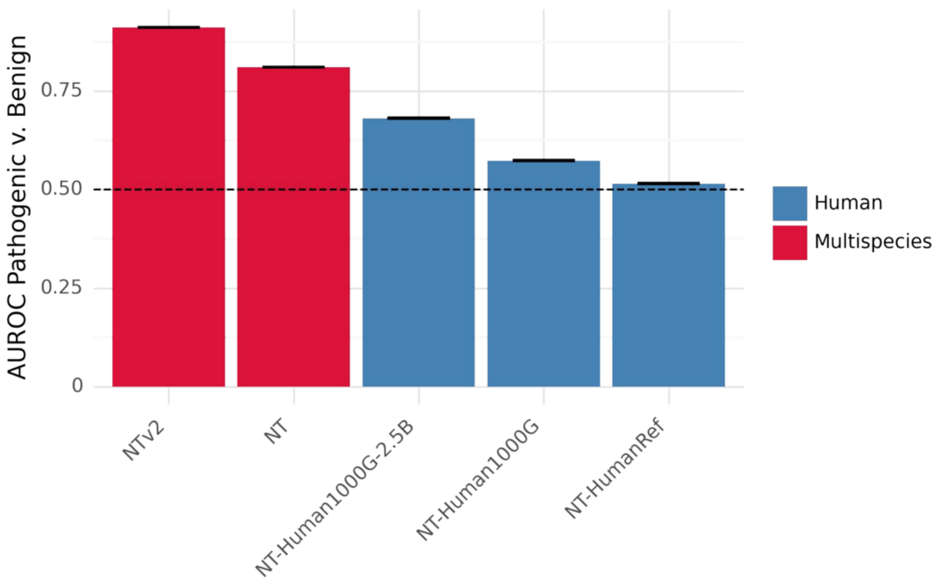


**Supplementary Figure 7. A)** Area under the ROC curve for absolute variant effect prediction on the same dataset as Fig. 1C using variant influence scores computed from Nucleotide Transformer models.

### Supplementary Table S1

| Model | Context | Resolution | Training Data | Architecture | Specialized for | Used in |
|---|---|---|---|---|---|---|
| SpeciesLM Fungi | 1 kb | Single bp | Fungi regions 5' of start codons (806 species) | Transformer | Promoters, 5'UTR, quasi Genome-wide* | Fig 1D, S1D,E,G,H ,2A,C,D,E 3B,C, S3A, S4A,B,C,5 A,B,C,D, S5A, 6A,B,C,D,7 A,B |
| SpeciesLM Metazoa | 2 kb | Single bp (overlapping 6-mer) | Metazoa regions 5' of start codons (494 species) | Transformer | Promoters, 5'UTR | Fig 1C, S1A,B,C,F 2B, 3A |
| SpliceBERT | 512 bp | Single bp | Metazoa pre-mRNA | Transformer | Splicing | Fig 3D,E |
| RiNALMo | 1 kb | Single bp | noncoding RNA | Transformer | noncoding RNA | Fig 4A,B,C,D,E ,F,G, 7A,B |
| Nucleotide Transformer NTv2 | 12 kb | 6-mer | Multispecies Genomes (Metazoa, Fungi, Prokaryotes) | Transformer | Genome-wide | Fig 7A,B,C, S7A |
| Nucleotide Transformer NT | 6 kb | 6-mer | Multispecies Genomes (Metazoa, Fungi, Prokaryotes) | Transformer | Genome-wide | Fig 7B,C, S7A |
| Nucleotide Transformer NT-Human1000G-2.5B | 6 kb | 6-mer | 3,202 diverse Human genomes | Transformer | Genome-wide | Fig 7A,B,C, S7A |
| Nucleotide Transformer NT-Human1000G | 6 kb | 6-mer | 3,202 diverse Human genomes | Transformer | Genome-wide | Fig 7B,C, S7A |
| Nucleotide Transformer NT-HumanRef | 6 kb | 6-mer | Human Reference Genome | Transformer | Genome-wide | Fig 7A,B,C, S7A |
| DNA-BERT | 512 b | Single bp (overlapping 6-mer) | Human Reference Genome | Transformer | Genome-wide | Fig 7B |
| Hyena | 1 Mb** | Single bp | Human | Hyena | Genome-wide | Fig 7A,B |

| | | | | | | |
|---|---|---|---|---|---|---|
| | | | Reference Genome | (Autoregressive) | | |
| Evo | 8 kb** | Single bp | Prokaryotic Genomes | Striped Hyena (Autoregressive) | Genome-wide | Fig 7A,B |
| CaduceusPS | 131 kb | Single bp | Human Reference Genome | Bidirectional Mamba | Genome-wide | Fig 7A,B |
| PlantCaduceus | 512 b | Single bp | 16 Plant Genomes | Bidirectional Mamba | Genome-wide | Fig 7A,B |

* Because many fungal genomes are compact, taking the region 1 kb 5' of gene starts already covers a significant amount of the genome, including diverse features such as non-coding RNA, coding sequences, regulatory elements, long-terminal repeats and others. ** Several versions trained for different lengths exist.
